# Computational and experimental analyses of mitotic chromosome formation pathways in fission yeast

**DOI:** 10.1101/2020.10.15.341305

**Authors:** Tereza Gerguri, Xiao Fu, Yasutaka Kakui, Bhavin S. Khatri, Christopher Barrington, Paul A. Bates, Frank Uhlmann

## Abstract

Underlying higher order chromatin organization are Structural Maintenance of Chromosomes (SMC) complexes, large protein rings that entrap DNA. The molecular mechanism by which SMC complexes organize chromatin is as yet incompletely understood. Two prominent models posit that SMC complexes actively extrude DNA loops (loop extrusion), or that they sequentially entrap two DNAs that come into proximity by Brownian motion (diffusion capture). To explore the implications of these two mechanisms, we perform biophysical simulations of a 3.76 Mb-long chromatin chain, the size of the long *S. pombe* chromosome I left arm. On it, the SMC complex condensin is modeled to perform loop extrusion or diffusion capture. We then compare computational to experimental observations of mitotic chromosome formation. Both loop extrusion and diffusion capture can result in native-like contact probability distributions. In addition, the diffusion capture model more readily recapitulates mitotic chromosome axis shortening and chromatin density enrichment. Diffusion capture can also explain why mitotic chromatin shows reduced, as well as more anisotropic, movements, features that lack support from loop extrusion. The condensin distribution within mitotic chromosomes, visualized by stochastic optical reconstruction microscopy (STORM), shows clustering predicted from diffusion capture. Our results inform the evaluation of current models of mitotic chromosome formation.

## INTRODUCTION

Dynamic chromatin organization during interphase is crucial for the regulation of gene expression and other nuclear processes. In mitosis, chromatin compacts to give rise to well-defined X-shaped chromosomes, a prerequisite for their faithful segregation. At the basis of higher order chromatin organization lie Structural Maintenance of Chromosomes (SMC) complexes, large protein rings that have the ability to topologically entrap DNA (1–3). SMC rings include an ATPase, suggesting that energy is expended to organize chromatin or to regulate the process. During interphase, the major chromosomal SMC complex is the cohesin complex that establishes cohesion between the newly replicated sister chromatids. It does so by topologically entrapping the two sister DNAs. Cohesin also participates in organizing interphase chromatin into topologically associating domains (TADs). As cells progress towards mitosis, a second SMC complex, condensin, rises in importance. Condensin is enriched, or activated, on mitotic chromosomes to promote chromosome compaction. Without condensin, chromosomes fail to reach their mitotic shape and are unable to segregate, leaving behind anaphase bridges. The molecular mechanism by which SMC complexes organize chromatin has remained a matter of debate. Two prominent models postulate that SMC complexes actively extrude DNA loops (loop extrusion), or that they sequentially entrap two DNAs that come into proximity by Brownian motion (diffusion capture).

Both condensin and cohesin have, under certain conditions, been observed to extrude DNA loops *in vitro* (4–7). Once bound to DNA, these SMC complexes asymmetrically or symmetrically reel in DNA, thereby forming a DNA loop. The observations suggest that little ATP is hydrolyzed to rapidly move over long distances. Applied to chromatin, condensin has been proposed to similarly reel in chromatin until it reaches a neighboring condensin complex that is itself engaged in loop extrusion. This would lead to formation of a central protein scaffold from which DNA loops emerge, reflecting chromosome models based on cytological and early biochemical analyses (8,9). Simulations of this process taking place on human chromosomes have shown agreement with experimentally observed chromosome formation, chromosome axis establishment and sister chromatid resolution (10). A feature of the loop extrusion model is that condensin-mediated DNA contacts will always lie within one chromatin chain. However, whether condensin can indeed extrude loops on a chromatin substrate densely decorated by histones and other DNA binding proteins remains unclear.

An alternative mechanism by which condensin can contribute to chromosome formation is by stabilizing stochastic pairwise interactions between condensin binding sites (11). We refer to this mechanism as ‘diffusion capture’. A condensin complex that has topologically loaded onto DNA might be able to embrace a second DNA that comes into proximity by Brownian motion. This mechanism could be akin to cohesin’s ability to capture a second DNA, following its loading onto a first DNA (12). Alternatively, two condensin complexes that each embrace one DNA might engage with each other. A tendency of SMC complexes to form clusters on DNA *in vitro* (13–15) is consistent with the latter possibility. In the diffusion capture scenario, condensin establishes contacts both within chromosomes and between chromosomes, consistent with experimental observations in yeasts (16–18). Computational simulation of diffusion capture taking place on a small budding yeast chromosome has generated chromosome properties with a good fit to experimentally observed chromosome behavior (11). The simulations also revealed that the intrinsically higher likelihood of condensin to establish interactions within a chromatin chain, as compared to between two independently moving chains, is sufficient to achieve chromosome individualization. Whether diffusion capture suffices to govern the formation of larger chromosomes is not known.

In this study, we developed a coarse-grained Brownian dynamics simulation of a chromatin chain, the size of the long left arm of fission yeast chromosome I. We use these simulations to explore the consequences of loop extrusion and diffusion capture on chromosome formation. We compare predictions from both models to experimental observations in fission yeast. Both loop extrusion and diffusion capture result in chromosome formation and chromosome contact distributions similar to those observed *in vivo.* In addition, diffusion capture provides an efficient means to recapitulate condensin-dependent chromosome axis shortening and volume compaction, as well as experimentally observed chromatin mobility changes inside mitotic chromosomes. Finally, the localization of condensin within mitotic chromosomes using STORM imaging reveals condensin clusters that are predicted to arise from diffusion capture. We conclude that diffusion capture represents an appealing mechanism that we propose contributes to chromosome formation in fission yeast.

## MATERIALS AND METHODS

### *S. pombe* strains and culture

All the *S. pombe* strains used in this study are listed in Supplementary Table 1. To construct the Cut14-SNAP strain, the SNAP coding sequence (New England Biolabs) was cloned into a pFA-based fission yeast C-terminal tagging vector, then the C-terminus of the endogenous *cut14^+^* locus was fused to *SNAP* by PCR-based gene targeting (19). Strains were cultured in Edinburgh minimal medium (EMM) supplemented with 2% glucose and 3.75 g/L of L-glutamic acid as a nitrogen source. To arrest cells in mitosis, 5 μg/mL of thiamine was added to the EMM culture to repress Slp1 expression and incubated for 3 hours at 25 °C. For Cut14 depletion, cells were incubated for 90 minutes after the addition of 5 μg/mL thiamine at 25 °C to repress both Slp1 and Cut14 expression, and then 0.5 mM of the auxin 3-indoleacetic acid (IAA) was added to the culture to degrade Cut14 and incubated for another 90 minutes at 25 °C before cells were collected.

### Measurement of DNA volume and chromatin loci distance

Cells were fixed with 70% ethanol and then stained with 4’,6-diamidino-2-phenylindole (DAPI). Images were acquired as serial sections along the z axis on a DeltaVision microscope system (Applied Precision). To measure the DNA volume, all the images were deconvolved in SoftWoRx and then the voxels over an arbitrary DAPI signal intensity threshold were counted using the 3D objects counter in Fiji (20,21). Distance distribution data between chromatin loci was adopted from (22).

### Chromatin mobility tracking and mean square displacement (MSD) calculation

For chromatin mobility tracking, a single focal plane of live cells was imaged at 20 ms intervals using a custom-built spinning-disc confocal microscope system (Intelligent Imaging Innovations) (22,23). The movement of a fluorescent dot was automatically traced using Virus Tracker (https://github.com/djpbarry/CALM/wiki/Virus-Tracker). The weighted mean of the MSD was calculated using the @msdanalyzer Matlab class (24). Further details are described in (21).

### Anisotropy of motion determination from trajectories at short times

From a trajectory in 2D, we determined anisotropic motion where the diffusion constant is not the same in all directions and/or there are different constraints in one direction or another. In both cases the MSD along each direction will be different. In the case of a polymer like chromatin, there is in general no good frame of reference, since the local environment rearranges over time. Over long times the MSDs along two axes will therefore be the same. However, over short times the local environment will be relatively constant, and the MSDs along two axes will show a difference if there are anisotropic constraints. For this reason, we define time-dependent anisotropy *η* in the following way:

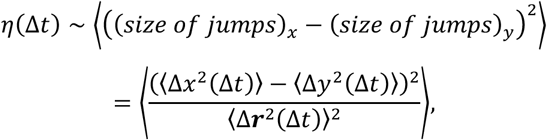

where (Δx^2^(Δt)) is the MSD in the x-direction, (Δy^2^(Δt)) is the MSD in the y-direction and (Δ**r**^2^(Δt)) = (Δx^2^(Δt)) +*{Δy^2^(Δt))* the total MSD in 2D. Note that *η* is an average over a number of trajectories, where for each trajectory the MSD is calculated by an average over all displacements with delays Δt. *η* is roughly the squared normalized average difference between the diffusion constants in *x* and *y* directions, *D_x_* and *D_y_,* and so we can roughly relate the ratio of these diffusion constants to *η* in the following way

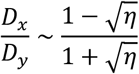

where without loss of generality we assume that *D_x_ ≤ D_y_,* by assuming the *x* direction is that corresponding to the smaller diffusion constant.

### STORM microscopy

Cells were fixed with 3.7% formaldehyde in PEM buffer (0.1 M PIPES, 1 mM ethylene glycol-bis (ß-aminoethyl ether)-N,N,N’,N’-tetraacetic acid (EGTA), 1 mM MgSO_4_) for 10 minutes at room temperature. Fixed cells were washed with PEM buffer containing 1.2 M Sorbitol three times. To permeabilize cell membranes, cells were treated with 0.1% Triton X-100 in PEM buffer for 5 minutes at room temperature. Cut14-SNAP was stained with 0.2 μM of SNAP-Surface Alexa Flour 647 (New England BioLabs) in PEM buffer for 15 minutes at 25 °C. After washing cells with PEM buffer three times, SNAP-stained cells were mounted on Nunc™ Lab-Tek™ II Chambered Coverglass 8 wells (Sigma) coated with Lectin. STORM imaging was performed in imaging buffer (20 mM Cysteamine (MEA, Sigma), 1% 2-mercaptoethanol (Sigma), 50 mM Tris-HCl (pH 8.0), 10 mM NaCl, 10% glucose, 205.4 U/mL Glucose Oxidase (Sigma), 5472 U/mL Catalase (Sigma)).

STORM images were collected on a Bruker Vutara 352 commercial 3D biplane single molecule localization microscope using a 60x silicone objective (Olympus) with a numerical aperture of 1.2 (25). We used a 640 nm laser with 50% laser power for illuminating Alexa Fluor 647 and a 405 nm laser with 0.5% laser power for photo-activation. Fluorescent signals were captured on an ORCA-Flash4.0 CMOS camera (Hamamatsu) using 20 ms exposure. We collected 30,000 frames and eliminated the first 10,000 frames for data processing.

To determine precise particle localization, we followed a previously described data processing method (25) with slight modifications. Briefly, we removed localizations with lower quality score (< 0.8, the value ranging from 0 to 1) according to the goodness-of-fit metric of each localization event. We then removed localizations that did not blink for longer than 3 frames. Finally, we eliminated all localizations with a lower axial precision (> 100 nm). Filtering was performed using Bruker’s SRX software.

### Simulation of a coarse-grained chromatin chain

A virtual chromatin chain was constructed to study the expected behavior of the long left arm of *S. pombe* chromosome I. The chain comprises 1,880 consecutively connected beads with a radius of 25 nm, each reflecting a string of ~10 nucleosomes covering a genomic size of ~2 kb. This chain thus corresponds to a genomic length of ~3.76 Mb, equivalent to the long *S. pombe* chromosome I left arm. Any two connected beads elastically interact and any two beads that overlap mutually repel. In the absence of introduced condensation mechanisms, each chromatin bead undergoes Brownian motion, constrained by attractive and volume exclusion forces. Effectively, the relaxed chain behaves as a self-avoiding Rouse polymer (Figure 1A).

**Figure 1.**
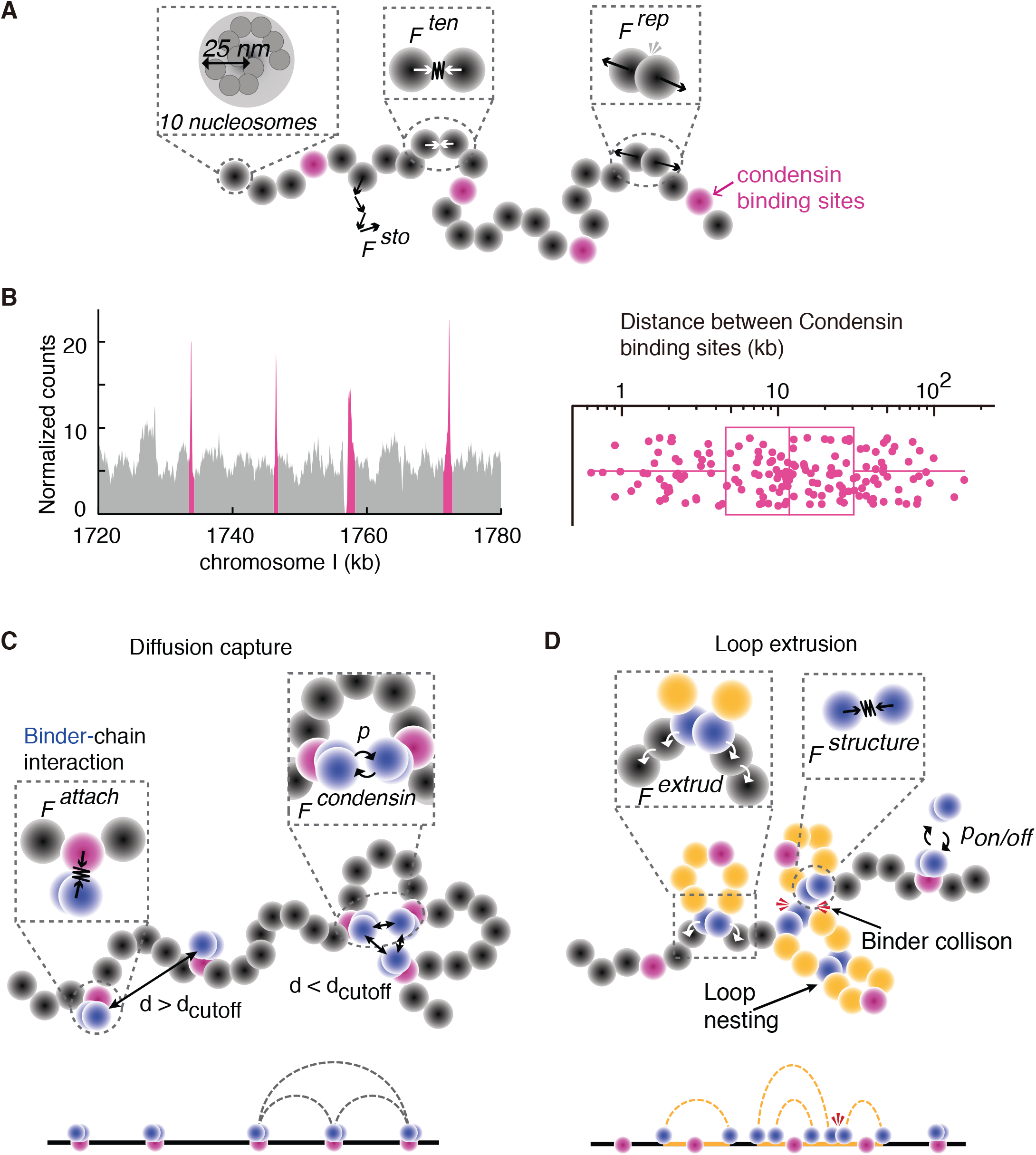
A biophysical model of the fission yeast chromosome I left arm. (**A**) Schematic of the coarse-grained chromatin polymer model and the forces exerted on the chromatin chain. Grey beads with a radius of 25 nm are equivalent of ~10 nucleosomes and represent a ~2.0 kb chromatin region. Condensin binding sites are highlighted in magenta. A stochastic force *(F^sto^)* allows each bead to follow a Brownian dynamic trajectory. The tension force *(F^ten^)* connects beads and constrains their movement, whereas a repulsion force *(F^rep^)* limits bead overlap. (**B**) Example of condensin localization along a 60 kb region in the middle of the chromosome I left arm (left; (22)). Condensin enriched sites are highlighted in magenta. The distance distribution between neighboring condensin binding sites along the chromosome I left arm are plotted (right), the box shows the median, 25^th^ and 75^th^ percentile, the whiskers indicate the range. (**C**) Schematic of the applied physical forces in the diffusion capture model. Condensin-chromatin association is secured by an attachment force *(F^attach^).* Two condensins are allowed to be attracted by a condensin capture force *(F^condensin^)* and form a diffusion capture pair if their Euclidean distance d < d_cutoff_. *F^condensin^* is additionally regulated by an association probability *p*. (**D**) Schematic of the forces in the loop extrusion model. Each condensin consists of two ‘feet’ that move in opposite directions. Movement is secured by the extrusion force *(F^extrud^)* that replaces *F^attach^* and targets beads one removed from the bead of residence. The two feet are prevented from splitting by a structure force *(F^structure^).* At certain time intervals, an association probably *p*_on/off_ allows condensin to detach and re-load onto a free condensin binding site to initiate a new loop or loop nesting. The resulting looping patterns in (**C**) and (**D**) are schematically illustrated.

#### Chromatin bead unit

A linear array of 10 nucleosomes with 10 nm diameter including linker DNA reaches just over 100 nm (radius 50 nm). Tight hexagonal packing of 10 nucleosomes in turn results in an assembly with radius 15 nm. This gives us upper and lower bounds for the size of a 10 nucleosome unit. Based on finegrained simulations of a histone chain (11), we observe that 10 nucleosomes in a chromatin chain typically occupy a volume with a radius of approximately 25 nm. This volume is only partially filled with nucleosomes and is accordingly modeled as a soft sphere without a rigid boundary.

#### Special sites on the chromatin chain

While the virtual chromatin chain is a homopolymer in a physical sense, a few beads are marked as special sites corresponding to their biological roles. The first and last beads of the chain represent the telomere and centromere, respectively. A group of beads with 0.1, 0.7, 1.2, 1.7, and 2.2 Mb genomic distance from a locus close to the centromere are labeled to correspond to fluorophore-tagged sites, allowing inter-fluorophore distances to be monitored akin to experimental observations (22,26). 158 beads are selected to be ‘condensin binding sites’. Their distribution is based on a condensin ChIP experiment in fission yeast (22). The mean distance between neighboring condensin binding sites is 11.7 beads (23.4 kb), the median distance is 6 beads (12 kb) (Figure 1B). These beads are either the ‘host’ sites of condensin to mediate diffusion capture or the starting positions of condensin to initiate loop extrusion.

#### Boundary condition and initial configuration

To resemble conditions in the interphase *S. pombe* nucleus, the chromatin chain was placed in a spherical volume of 14.14 μm^3^ (1.5 μm radius) with a rigid boundary to represent the *S. pombe* nucleus. *S. pombe* interphase chromatin contains few defined structural domains, such as TADs (22,27). For this reason, a set of ‘random’ conformations was created within a cylindrical subsection (3.84 μm^3^) of our virtual nucleus, corresponding to the fraction that the chromosome I left arm represents of the total fission yeast genome. The cylindrical constraint was removed and evolution of the initialized chromatin chain was then subject to the rules and physics-based forces introduced below.

#### Forces employed

In the absence of active processes governing condensation, a bead *i* in the chromatin chain is subject to a stochastic force 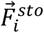 sourced from collision with molecules in the nucleoplasm, a tension force 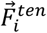 imposed by connected beads, and a volume-exclusion repulsive force 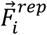 exerted by spatially overlapping beads. Additionally, a damping force 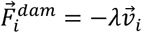 acts on the moving bead, representing the viscous effect of the nucleoplasm, the magnitude of which is assumed to be proportional to instantaneous speed.

##### 1. Stochastic force

A stochastic force is applied to each bead, both chromatin beads and condensins (see below), at each simulation step as:

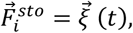

Any component of the stochastic force 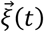 has a time-average of zero and is uncorrelated in space and time. Namely,

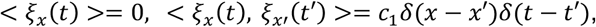

In practice, at each step, the instantaneous value of any component of the force 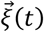 is calculated as 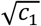 multiplied by a random number drawn from a Gaussian distribution with a zero mean and a standard deviation of 1. The constant c_1_ is set as:

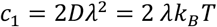

with Stokes-Einstein relation:

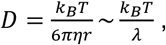

Where *D* represents the diffusion coefficient, *η* dynamic viscosity and *r* is the bead radius. Since the coarse-grained bead does not represent a rigid sphere but rather a flexible chain of ~10 nucleosomes, the relation *λ = 6πηr* does not apply. For simplicity, we introduced a plausible damping constant *λ.*The value of c_1_ allows the bead to have an average movement on a scale consistent with experimental observations (21,28). Coarse-grained bead movement is principally regulated by the entropic force and the spring constant of the chromatin bead linker. This parameter pair was chosen such that the bead displacement distribution over short (20 ms) time intervals was compatible with that observed in the *S. pombe* interphase nucleus (22).

##### 2. Tension force

A linear elastic force (e.g. Hookean spring) is applied to describe the interaction between two consecutively connected coarse-grained beads:

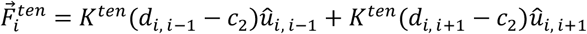

where *K^ten^* is the spring constant of the linker, *d_i, i+1_* is the distance between the centers of two consecutively connected beads *i* and *i* + 1; c_2_ is a constant describing the equilibrium (non-stretched or non-compressed) length of the bead linker; *û_i, i+1_* are unit vectors determining the direction of the force.

##### 3. Repulsion force

In order to limit overlaps between any two beads, a constant volume exclusion force between two beads within *d_rep0_* < 50 nm of each other is applied. Unless stated otherwise, *û_a_, _b_* denotes a unit vector from object *a* to object *b*.

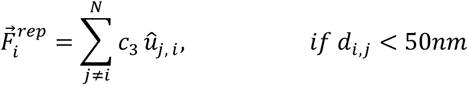

where *c*_3_ is a constant equal to 0.5; j is a bead different from i.

### Diffusion capture simulations

The diffusion capture model describes the crosslinking of distal genomic sites mediated by condensin. To implement this model an additional pair of beads, representing a condensin molecule, is bound to condensin binding sites. When two condensins bound to different binding sites stochastically become spatially adjacent, they have a probabilistic propensity of association (Figure 1C). In some simulations, we titrated the condensin concentrations such that we removed randomly 85, 75, 50 or 25% of condensin molecules from their binding sites.

#### Forces employed

Three additional forces are employed in order to implement the diffusion capture model. A *condensin structure force* describes the interaction between the two condensin ‘feet’ that reflects the structural integrity of a condensin molecule; a *condensin attachment force* describes the interaction between both condensin feet and a chromatin bead that maintains condensin attachment to the chromatin chain; a *condensin capture force* describes the interaction between condensins on different beads that mediates diffusion capture.

##### 1. Condensin structure force

While condensin is modelled as two beads, a front and a rear ‘foot’, only the front foot participates in diffusion capture. A linear elastic force is applied between the two feet to maintain their spatial proximity, which becomes important later in the loop extrusion model.

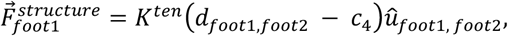

where c_4_, representing the equilibrium distance between the two feet, equals 0 nm. The radius of both feet is 25 nm.

##### 2. Condensin attachment force

The interaction between each condensin foot and its binding site is described as:

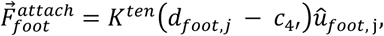

where *c_4_,* is equal to 0 nm; *j* refers to the chromatin bead that condensin is attached to.

##### 3. Condensin capture force

The condensin capture force, 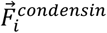 is applied between condensin front feet on different binding sites as an elastic spring following Hooke’s law:

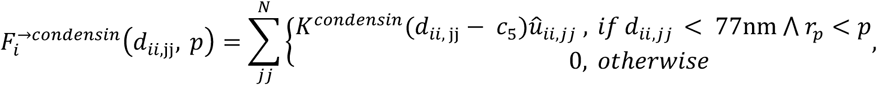

This force is exerted when the center of front foot *ü* and the center of another front foot *jj* are within a cut-off distance *d*_*ii*, jj_ =*77nm.* This equates to a distance of 27 nm between the bead surfaces, a conservative estimate for a distance that might be bridged by a condensin molecule. *p* is the dissociation probability which represents turnover of diffusion capture pairs. Algorithmically, it is implemented through a random number generated at each time step for each 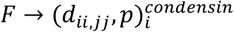 if a random number *r_p_* is less than a threshold *p* then 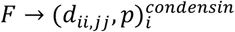 becomes zero. *c_5_* is the equilibrium distance between two interacting condensins. Here, we define *c_5_* as 52 nm, meaning that two condensins lie adjacent. The valence of diffusion capture sites, 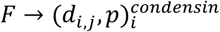 representing the number of interacting partners, is naturally regulated by the forces in the system and steric constraints.

### Loop extrusion simulations

In contrast to diffusion capture, where condensins attach to their binding sites and remain in position, in the loop extrusion model condensins load at empty binding sites from where they translocate. The two condensin feet symmetrically move in opposite directions along the chromatin chain by repeatedly associating with the next chromatin bead, thereby bridging distant genomic sites to form a chromosome loop (Figure 1D). When two condensin complexes encounter each other, movement of colliding feet is stopped. Condensin feet that are not in collision continue translocation, resulting in further asymmetric loop extrusion. The rate of translocation is given by:

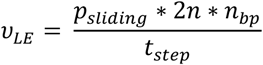

Where *p_sliding_* is a probability of translocation, *n* is the number of travelled beads, *n_bp_* represents the DNA length in bp per bead and *t_step_* is the simulation timestep. In our simulation, *ν_LE_ ~* 1.2 kb/s in line with experimentally observed values (4,29). To introduce condensin turnover, each condensin can stochastically unbind from the chromatin chain and relocate. A constant condensin concentration on chromatin is achieved such that every time a condensin is unloaded, a new condensin is loaded at an empty binding site. Algorithmically, dynamical condensin exchange is implemented such that at *T_exchange_* time intervals a dissociation probability *p_on/off_* is calculated (akin to the condensin dissociation probability *p* in the loop extrusion model) for each condensin to decide whether it is unloaded from its current position and relocated to an empty binding site.

### Forces employed

The loop extrusion model differs from diffusion capture in that the *condensin attachment force* is repurposed as an *extrusion force* 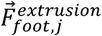, acting to elastically connect the translocating condensin foot with the associated chromatin bead.

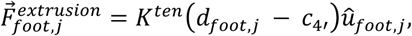

where 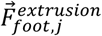 is a tension force allowing a condensin foot to interact with bead *j* on the chromatin chain. Bead *j* is iteratively being updated to the following chromatin bead *j* + 1 (in the case of forward-moving condensing foot) or *j* – 1 (in the case of backward-moving condensing foot), therefore allowing translocation of condensin along the chromatin chain and re-assignment of their 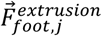 from the current chromatin bead to the adjacent one. A *condensin capture force* is not operational in the loop extrusion model.

### Dynamics and model implementation

The overdamped Langevin equation is employed to describe the time evolution of the coarse-grained chromatin configuration. This assumes that the inertial part 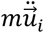 is much smaller than the damping part 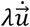, on the longer timescales of interest in this study. Under this assumption, the dynamical equation to describe a free chromatin chain is as follows:

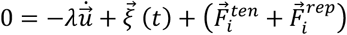

or

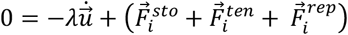

In the diffusion capture model, additional forces are included to describe the movement of a specific bead *i*:

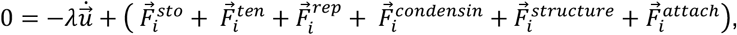

In the loop extrusion model, movement of a specific bead *i* during simulation is controlled by a summation of forces:

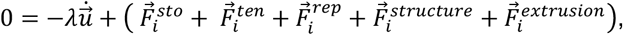

At each simulation step, the Euler integration has been applied to the dynamics equation in order to describe time evolution of the system, therefore movement of each bead is described for velocity *v_ix_(t)* and tension *u_ix_(t*+*Δt)* as follows:

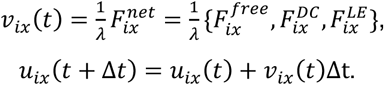

Where *F^free^, F^DC^, F^LE^*, as given by the sum of terms in the equations above, correspond to forces employed in free chromatin chain, the diffusion capture model, and the loop extrusion model, respectively.

### List of parameters regulating bead movement

| Parameters | Values | Dimension | Host function |
| --- | --- | --- | --- |
| $\lambda$ | $3 \times 10^{-8}$ | $kg/s$ | $F_i^{\rightarrow sto}$ |
| $K^{ten}$ | $1 \times 10^{-1}$ | $pN/nm$ | $F_i^{\rightarrow sto}$ |
| $d_{i,i+1}$ | *** | $nm$ | $F_i^{\rightarrow sto}, F_i^{\rightarrow ten}, F_i^{\rightarrow rep}$ |
| $c_2$ | $5.2 \times 10^2$ | $nm$ | $F_i^{\rightarrow ten}$ |
| $u_{i,i+1}^{\wedge}$ | *** | - | $F_i^{\rightarrow sto}, F_i^{\rightarrow ten}, F_i^{\rightarrow rep}$ |
| $c_3$ | $5 \times 10^{-1}$ | $pN$ | $F_i^{\rightarrow rep}$ |
| $d_{rep0}$ | $5 \times 10^1$ | $nm$ | $F_i^{\rightarrow rep}$ |
| $K^{binder}$ | $1 \times 10^{-1}$ | $pN/nm$ | $F_i^{\rightarrow binder}$ |
| $c_4$ | 0 | $nm$ | $F_i^{\rightarrow binder}$ |
| $K^{ten\_binder}$ | $1 \times 10^{-1}$ | $pN/nm$ | $F_i^{\rightarrow attach}$ |
| $c_{4'}$ | 0 | $nm$ | $F_i^{\rightarrow attach}$ |
| $K^{condensin}$ | $1 \times 10^{-1}$ | $pN/nm$ | $F_i^{\rightarrow condensin}$ |
| $c_5$ | $5.2 \times 10^2$ | $nm$ | $F_i^{\rightarrow condensin}$ |
| $d_{condensin0}$ | $7.7 \times 10^2$ | $nm$ | $F_i^{\rightarrow condensin}$ |
| $p_o$ | $1 \times 10^{-2}$ | - | $F_i^{\rightarrow condensin}$ |
| $d_{i,j}$ | **** | $nm$ | $F_i^{\rightarrow structure}$ |
| $u_{i,j}^{\wedge}$ | **** | - | $F_i^{\rightarrow structure}$ |
| $d_{ii,jj}$ | **** | $nm$ | $F_i^{\rightarrow condensin}, F_i^{\rightarrow attach}$ |
| $u_{ii,jj}^{\wedge}$ | **** | - | $F_i^{\rightarrow condensin}, F_i^{\rightarrow attach}$ |
| $T_{exchange}$ | 2 | $s$ | condensin turnover during LE |
| $p_{on/off}$ | $1 \times 10^{-2}$ | | condensin turnover during LE |

### Measurements and readouts

All simulations were run for 1,200 seconds with a simulation timestep ***dt* = 10^−4^** s. Each simulation condition for diffusion capture and loop extrusion was recapitulated with 10 simulation replicates. If not stated otherwise, readouts were collected every 10 seconds, resulting in 1200 measurements from the 10 replicates.

#### Computational fluorophore distance measurements

We mapped experimental fluorophore arrays (22,26) onto the computational chromatin chain and measured Euclidean distances between selected fluorophore pairs.

#### Computational Hi-C and interaction frequency analysis

We generated Hi-C-like representations of chromosome conformations during our simulations. Instead of contact frequency, we display Euclidean distance between any two chromatin beads, averaged over 12,000 conformations collected at one second intervals from the 10 simulation replicates. To plot interaction frequency as a function of genomic distance, we considered two beads as interacting if their Euclidean distance was within 500 nm. The principle conclusions from this analysis were insensitive to the chosen cutoff. All bead pairs were binned according to their genomic separation to generate a frequency distribution. The frequencies are normalized to have a sum of one across all bins.

#### Computational volume measurements

To facilitate volume measurements, we divided our system into 3D voxels (cubes). Each voxel has a dimension of 100 nm. We define the volume of the chromatin chain as the total volume of occupied voxels.

#### Condensin clustering analysis

Clustering is performed using a DBSCAN algorithm in the open-source python library sklearn.cluster. A cutoff distance of 100 nm between centers of the two feet of individual condensins is selected to reflect two condensin diameters in the model. A minimum number of condensins per cluster of 2 is chosen for the comparison between STORM data and the computational diffusion capture and loop extrusion models.

#### Simulation MSD measurements

In order to determine the MSD exponent of chromatin mobility in our simulations, we used the same approach as for the experimental data. Instead of the experimental fluorophore labelled chromatin locus, we tracked the position of the 225 condensin binding sites. We analyzed the MSD exponent for each 2 second window, collected every 60th second during the simulations. Since in the experiments we only observe a randomly oriented 2D projection of the full 3D fluorophore motion, we applied a 3D to 2D projection of particles in our simulations. We found previously that this projection does not alter the mean MSD exponent, but slightly broadens the distribution (21).

#### Simulation anisotropy measurements

Anisotropy of chromatin bead motion in our simulations was determined as described for the experimental data during the same time windows as the MSD exponents.

## RESULTS

### A biophysical model of diffusion capture and loop extrusion along the fission yeast chromosome I left arm

To study fission yeast chromosome condensation, we developed a biophysical model of a chromatin chain representing the length of the long left arm of fission yeast chromosome I. Our coarse-grained chromatin chain consists of 1,880 beads, each covering a ~2 kb region corresponding to ~10 nucleosomes, representing 3.76 Mb of genomic distance (Figure 1A). A stochastic force *(F^sto^)* is applied to every bead of the system, under the assumption that a chromatin bead follows Brownian motion in isolation. Any two consecutive beads interact via a spring-associated tension force (*F^ten^*) following Hooke’s law. This results in collective dynamic behavior of a joined chromatin chain. In addition, a repulsion term is employed when beads overlap *(F^rep^),* taking into account the soft nature of the chromatin chain within each bead. We consider the behavior based on *F^sto^, F^en^* and *F^rep^* to be that of a free chromatin chain (Figure 1A). To simulate diffusion capture and loop extrusion, we selected specific beads along this polymer chain as condensin binding sites, recapitulating the experimentally observed condensin distribution along fission yeast chromosome I (Figure 1B) (22). These condensin binding sites are the ‘host’ sites of condensin to mediate diffusion capture, or the starting positions for condensin to initiate loop extrusion.

Condensin is modeled to comprise two ‘feet’ that are initially concentric with each other and the condensin binding site. Only the ‘front’ foot takes part in diffusion capture, the ‘rear’ foot gains relevance during loop extrusion. To model diffusion capture, condensin is attached to the chromatin bead via a spring-based attachment force *(F^attach^)* and remains bound to the same bead throughout the simulation. If two condensins on distinct chromatin beads encounter each other by stochastic movements they form a pairwise interaction with a defined probability via a condensin capture force *(F^condensin^,* Figure 1C). When multiple condensins spatially meet at a common place, they are able to form larger clusters, limited in size only by the geometric constraints of the system. The dynamic nature of diffusion capture is regulated by the association probability, which not only controls formation of new diffusion capture pairs, but also their maintenance at every simulation step.

In the loop extrusion model, the condensin attachment force is repurposed as an extrusion force *(F^extrusion^).* Condensin initially binds to a condensin binding site, from where its front and rear feet start translocating into opposite directions. *F^extrusion^* sequentially targets chromatin beads next to the current bead of residence, resulting in symmetric loop extrusion (Figure 1D). The two condensin feet remain connected to each other by a condensin structure force *(F^structure^).* When two condensins encounter each other, movement of the colliding feet is stopped, while feet that are not in collision continue translocation, resulting in asymmetric loop extrusion until they also encounter another condensin. Loop extruding condensins periodically have a chance to unload and load again at a free condensin binding site, thus ensuring dynamic loop formation and loop nesting. Parameters are chosen to match experimentally observed loop extrusion rates (4,29).

Fission yeast condensin accumulates in the nucleus in mitosis. During interphase, nucleo-cytoplasmic shuttling leads to condensin redistribution and equalization between the compartments (30,31). We have previously determined the nucleus-to-cytoplasm ratio in fission yeast to be 0.14 ± 0.05 (21). With therefore approximately 15% of nuclear condensin, we use 15% occupied condensin binding sites to represent *in silico* interphase, while we refer to 100% condensin binding site occupancy as *in silico* mitosis. Further details on the computational implementation of the diffusion capture and loop extrusion models can be found in the Materials and methods.

### Axial chromosome compaction by diffusion capture and loop extrusion

Axial shortening is a hallmark of condensin-dependent mitotic chromosome formation in yeasts (22,26,32–34). To inspect axial chromosome compaction, we monitored the distance of two fluorophore-marked loci at 1.78 Mb distance from each other *in vivo* and of similarly spaced *in silico*-marked loci in our model. The median *in vivo* interphase distance, projected onto a 2D plane, was 1.1 μm in interphase, which shortened by ~ 39% to 0.65 μm in mitosis (Figure 2A) (22). The distance of the same fluorophore pair was previously measured in 3D to around 1.8 μm in interphase contracting to around 1.0 μm (i.e. by 44%), in mitosis (26). Mitotic compaction in both studies depended on condensin.

**Figure 2.**
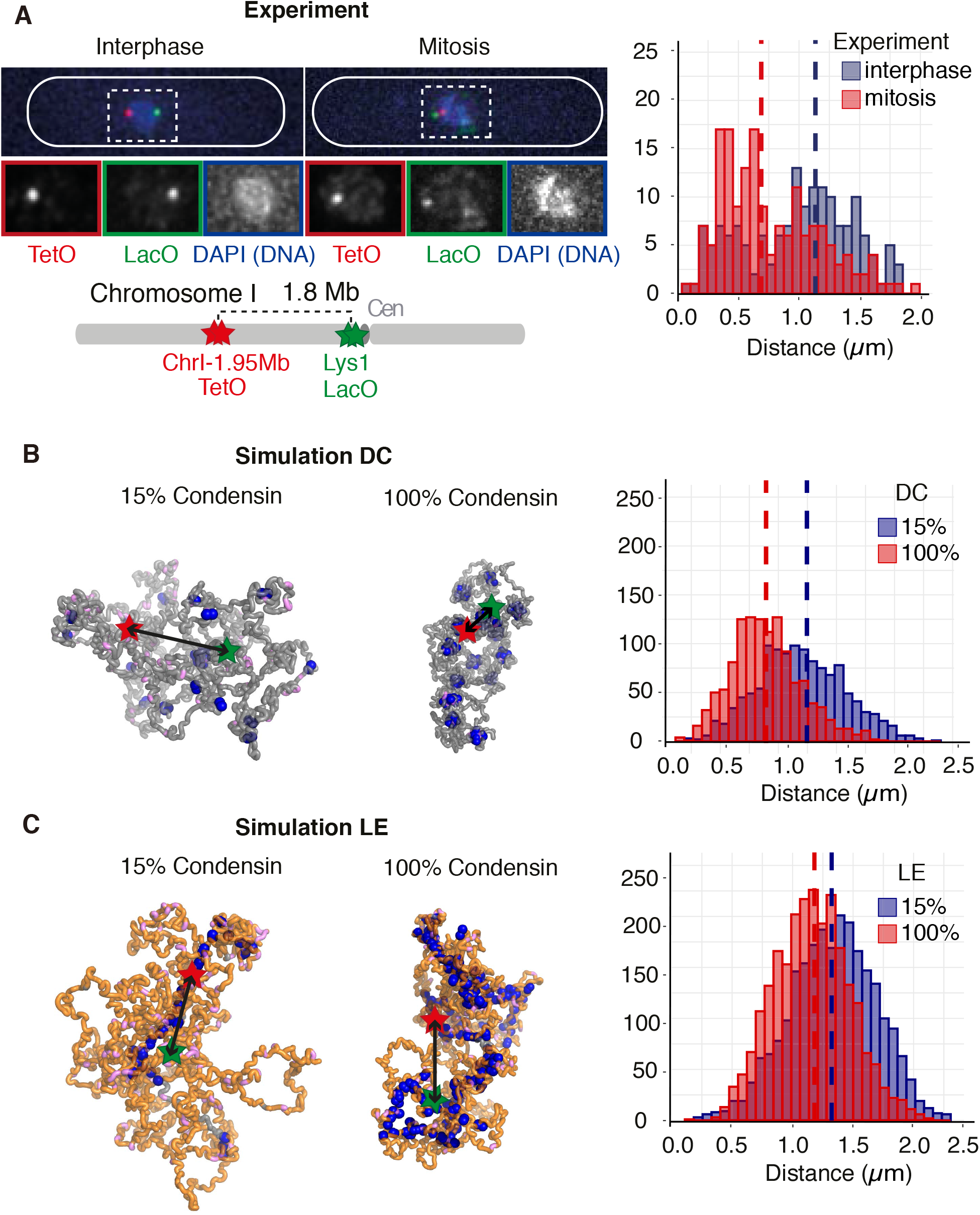
Axial chromosome compaction by diffusion capture and loop extrusion. (**A**) Fluorescent microscopy images of interphase and mitotic cells, showing two genomic loci marked by TetOs (red) and LacOs (green) together with DAPI staining of the DNA (blue). The schematic depicts the positions of the two loci along the chromosome I left arm (26). Distance distributions and their medians from >162 cells in each condition are shown. (**B, C**) Representative snapshots of *in silico* chromosome conformations by the diffusion capture (**B**) and loop extrusion models (**C**). Examples are shown of *in silico* interphase (15% condensin) and mitosis (100% condensin). Red and green stars represent *in silico* fluorophores, corresponding to those in (**A**). Their physical distance distribution and medians from 1200 snapshots of 10 simulation replicates is shown.

We started our computational simulations of diffusion capture from a relaxed chromatin chain, with either 15% (interphase) or 100% (mitosis) of occupied condensin binding sites. Diffusion capture pairs start to form and the system approaches a steady state when the number of capture pairs and the *in silico* fluorophore distance fluctuates around a constant value (Supplementary Figure S1A). Figure 2B shows representative conformations of our computational chromosome in both conditions. The real time movements of the chromatin chain can be observed in Supplementary Movies S1 and S2, illustrating frequent exchange of diffusion capture pairs in the steady state. We recorded 1200 3D fluorophore distance measurements at regular time intervals from 10 independent simulation repeats. These measurements show a well-defined distribution with a median of 1.2 μm in interphase and 0.89 μm in mitosis, roughly compatible with experimentally observed 3D distances and corresponding to a 26% mitotic chromosome axis shortening due to diffusion capture.

We next turned to the loop extrusion model. Upon the initiation of loop extrusion using either 15% or 100% of condensin per loading site, loops rapidly form and an axial condensin accumulation becomes discernable over time (Supplementary Figure S1B and Supplementary Movies S3, S4). At the interphase condensin concentration, a relatively short axial structure forms with long chromatin loops (Figure 2C). The *in silico* fluorophore distance is influenced by where the fluorophores find themselves relative to the axis, with a median distance of 1.3 μm in interphase. At the higher mitotic condensin concentration a greater number of loops, including a greater fraction of nested loops, are formed. This results in shorter loops and correspondingly a longer chromosome backbone. The fluorophore distance now depends on how the backbone arranges itself inside the chromosome, resulting in a simulated median Euclidean fluorophore distance of 1.2 μm. This corresponds to an 8 % chromosome arm shortening, less than what was achieved by diffusion capture.

To further explore the relationship between genomic and Euclidean distances in the diffusion capture and loop extrusion models, we inspected chromatin beads at 0.1,0.7, 1.2, 1.8 and 2.2 Mb distance, corresponding to previously experimentally observed fluorophore pairs (26). *In silico* interphase in either the diffusion capture or loop extrusion models recapitulated *in vivo* measured interphase distances reasonably well (Supplementary Figure S2A). Diffusion capture resulted in mitotic axial compaction in almost all observed cases, albeit not to the full extent that is observed *in vivo* (Supplementary Figure S2B). Thus diffusion capture makes a robust contribution to mitotic axial chromosome compaction. In contrast, loop extrusion often exhibited the opposite trend, generating increased mitotic Euclidean distances. In the loop extrusion model, the lengthening chromosome backbone due to additionally activated condensin appears to counteract axial compaction. Additional mechanisms might be required to achieve reproducible chromosome axis compaction in the loop extrusion model.

### *In silico* contact probability distributions due to diffusion capture or loop extrusion

Chromatin contact probability distributions, obtained from high throughput conformation capture (Hi-C) experiments, contain important information on chromosome architecture (35). Condensin enhances longer-range chromatin contacts during mitotic chromosome condensation at the expense of local chromatin contacts (22,36,37). Figure 3A shows experimental Hi-C maps of the fission yeast chromosome I left arm in interphase and mitosis, as well as the Hi-C interaction frequencies plotted as a function of their genomic distance. This illustrates enhanced mitotic chromatin interactions in a distance range from approximately 90 – 900 kb, which depend on condensin (22).

**Figure 3.**
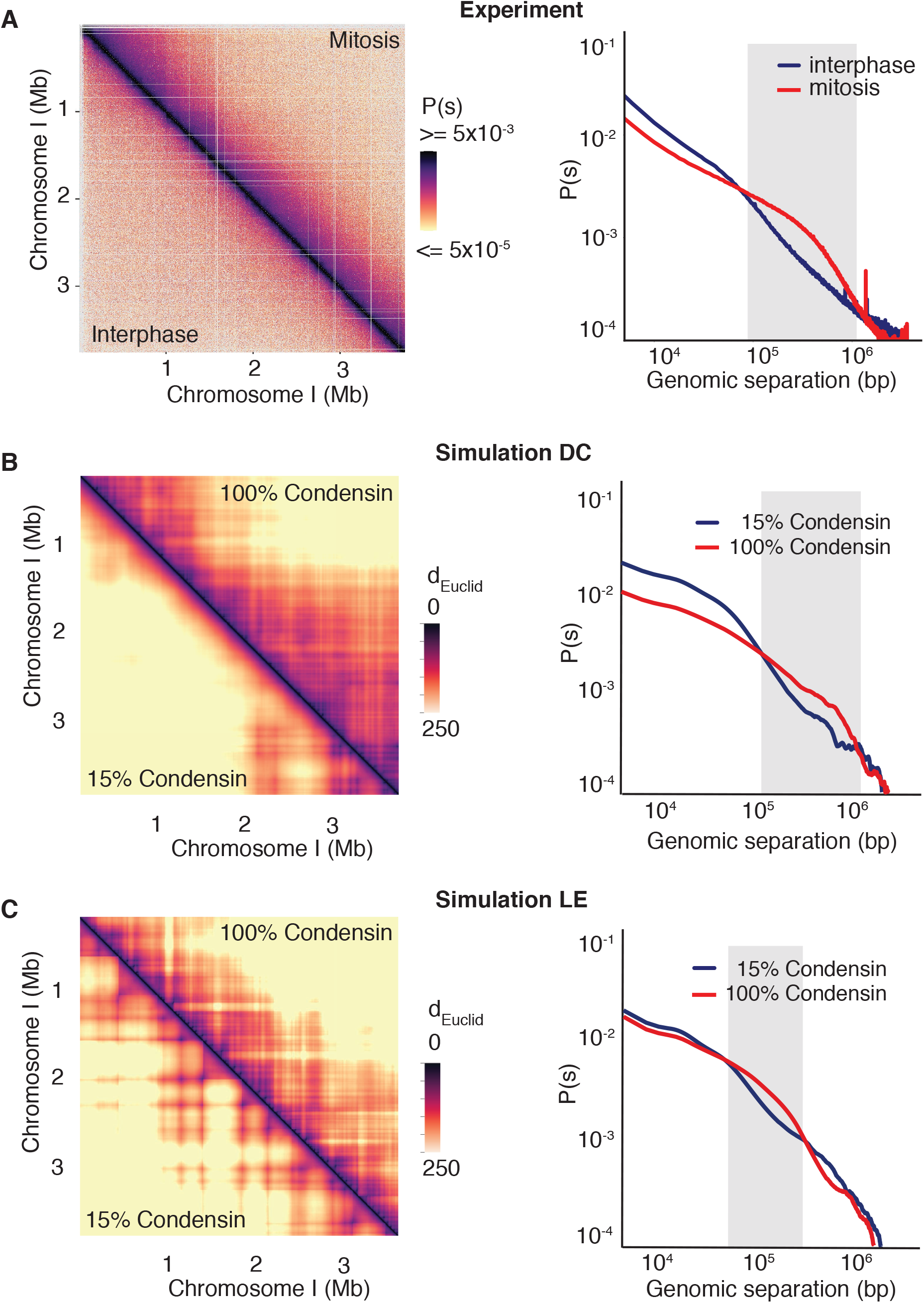
Contact probability distributions due to diffusion capture or loop extrusion. (**A**) Experimental Hi-C contact probability map of the fission yeast chromosome I left arm in interphase (lower triangle) and mitosis (upper triangle). The contact probability as a function of chromosomal distance along the chromosome arm is shown (22). The distance range of interactions that are augmented in mitosis (red), compared to interphase (blue) is shaded grey. (**B, C**) *In silico* Hi-C-like Euclidean distance maps averaged over 12,000 simulated chromosome conformations recorded in 1 second intervals during 10 simulation replicates under interphase and mitotic conditions in the diffusion capture (**B**) and loop extrusion models (**C**). Contact probabilities as a function of chromosomal distance are plotted using a 500 nm distance cut-off for scoring interactions. Regions where contacts are more frequent under mitotic conditions (red) compared to interphase (blue) are shaded grey.

To generate Hi-C-like depictions of our computational chromosome conformations, we display Euclidean distance maps, averaged over time and between simulation replicates (Figures 3B, C). These maps reveal that, in both the diffusion capture and loop extrusion models, the increased mitotic condensin concentration results in increased longer-range proximities, as seen by an expanded mitotic diagonal. To analyze interaction frequencies as a function of genomic distance, we set an arbitrary Euclidean distance cutoff at 500 nm to convert proximity into *‘in silico* Hi-C interactions’. In the case of diffusion capture, the interaction frequency plot reveals increased mitotic interactions over a distance range of 120 – 1,100 kb (Figure 3B), in approximate agreement with the experimental observations. Loop extrusion also resulted in increased longer-range interactions, albeit at a somewhat shorter distance range of 60 – 600 kb (Figure 3C). Thus, both *in silico* diffusion capture and loop extrusion recapitulate condensin-dependent mitotic chromatin contact changes.

To better understand the distance range of enhanced mitotic chromatin interactions, we titrated the condensin concentration in our simulations. In the case of diffusion capture, the interaction frequency plot of a free chromatin chain (0% condensin) showed only little difference from our interphase conditions (15% condensin). As soon as additional condensin binding sites were activated (25%), chromatin interactions in the 120 – 1100 kb distance range were augmented. Interactions increased further as more condensin binding sites were added, while their distance distribution remained roughly constant (Supplementary Figure S3A). Loop extrusion showed a different response pattern. Compared to the free chromatin chain, 15% of condensin resulted in an increase in chromatin interactions longer than 200 kb. This is likely explained by the formation of long chromatin loops in the presence of low condensin levels. As the condensin concentration increased, the distance range of chromatin interactions shortened, as expected from shortening chromatin loops. A closer match to the experimental interaction frequency distribution was obtained at intermediate condensin levels (Supplementary Figure S3B). Thus, both the diffusion capture and loop extrusion models reproduce experimental interaction frequency distributions. The distance range of enhanced mitotic interactions is robust in the case of diffusion capture, but sensitive to the condensin concentration in the case of loop extrusion.

### Chromatin volume compaction in mitotic chromosomes

A visually striking aspect of mitotic chromosome condensation is the volume reduction during the conversion of diffuse interphase chromatin into distinct chromosome bodies (38). In human cells this entails a ~2-fold volume compaction (39). Indeed, chromosome compaction was one of the first described roles of the fission yeast condensin complex (32). To quantify fission yeast chromosome compaction, we measured the chromatin volume in interphase and mitosis by 3D reconstructing serial z-stacks of fluorescent microscopy images of DNA stained with 4’,6-diamidino-2-phenylindole (DAPI). The median interphase chromosome volume was 2.06 μm^3^ which decreased in mitosis to 1.64 μm^3^, a 20% volume reduction (Figure 4A). Mitotic compaction depended on the condensin complex and was no longer observed following condensin depletion using a combined transcriptional shut-off and auxin-inducible degron strategy (40).

**Figure 4.**
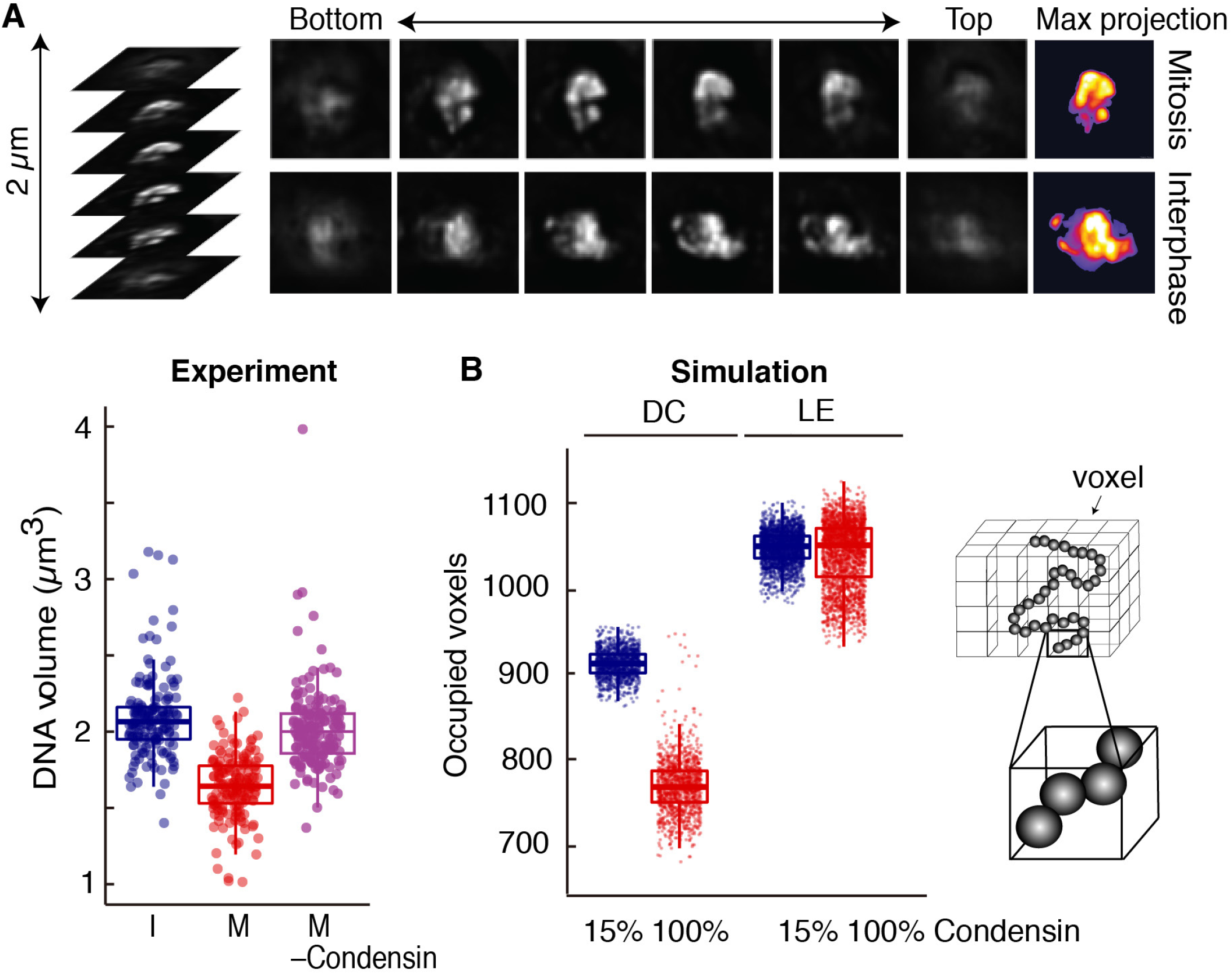
Chromatin volume compaction in mitotic chromosomes. (**A**) Examples of z-stacked images of DAPI-stained DNA in interphase and mitotic nuclei. Pseudocolor images of maximum intensity projections of the DNA volume are shown on the right. The volume distribution of >155 cells in interphase (I) and mitosis (M), as well as in mitosis following condensin shut-off (- Condensin) are shown. Boxes show the median and interquartile ranges. (**B**) Volumes of 1,200 simulated chromosome conformations, recorded every 10 seconds from 10 simulation replicates, as measured by occupied voxel counts using interphase and mitotic conditions during diffusion capture and loop extrusion simulations. Boxes depict medians and interquartile ranges. A schematic of chromatin beads, distributed across voxels, is included.

To measure chromatin volume in our simulations, we divided the nuclear volume into 100 nm-sized cubic voxels. We counted a voxel as occupied if it contained at least one chromatin bead. The chromosome I left arm accounts for approximately 20% of the fission yeast genome. Its *in silico* volumes were somewhat larger than the corresponding fraction of the experimentally measured DNA volume. This is likely the case because chromosomes lie close together in the yeast nucleus, reducing their apparent occupied volume at our microscopic resolution. Despite the different numerical values, the diffusion capture model resulted in an 16% volume reduction when comparing interphase and mitosis (Figure 4B). Volume reduction was condensin concentration-dependent (Supplementary Figure S4). This observation demonstrates that establishment of stochastic pairwise interactions between condensin binding sites along a chromatin chain can contribute to chromatin volume compaction. In contrast, loop extrusion resulted in only marginal volume changes. Specifically, the median volume increased by 0.09% during simulated mitosis (Figure 4B and Supplementary Figure 4). While interactions between distant parts of the genome are established by loop extrusion, the intervening chromatin is extruded, which limits the potential for volume compaction. Increased loop nesting, beyond that achieved based on simple probability, might be able to achieve increased levels of compaction in this model.

### Diffusion capture results in mitotic chromatin mobility reduction

During mitotic chromosome formation, condensin imposes constraint on the free movement of the chromatin chain (22). To experimentally study chromatin movements, we track a chromatin locus in the middle of the chromosome I left arm, marked by tandem lac operators bound by a LacI-GFP fusion protein. We then plot its mean squared displacement (MSD) over time. During interphase, we find that the MSD exponent over short time intervals is 0.49 ± 0.02 (mean ± 95% confidence interval, n = 595), consistent with a polymer chain whose diffusive behavior is only slightly constrained by a small amount of condensin (Figure 5A) (21). In mitosis, the exponent is markedly reduced to 0.28 ± 0.02 (n = 271). Looking more carefully at the distribution of MSD exponents from individual chromatin traces, the interphase distribution is well described by a single Gaussian fit. The mitotic sample, in contrast, showed a bimodal distribution that likely arose from contamination with a small number of interphase cells. A pure mitotic MSD exponent might be lower than 0.28, possibly as low as 0.23 (Supplementary Figure S5A). Exemplar trajectories of the GFP-marked locus over time illustrate the reduced mitotic chromatin mobility (Figure 5A).

**Figure 5.**
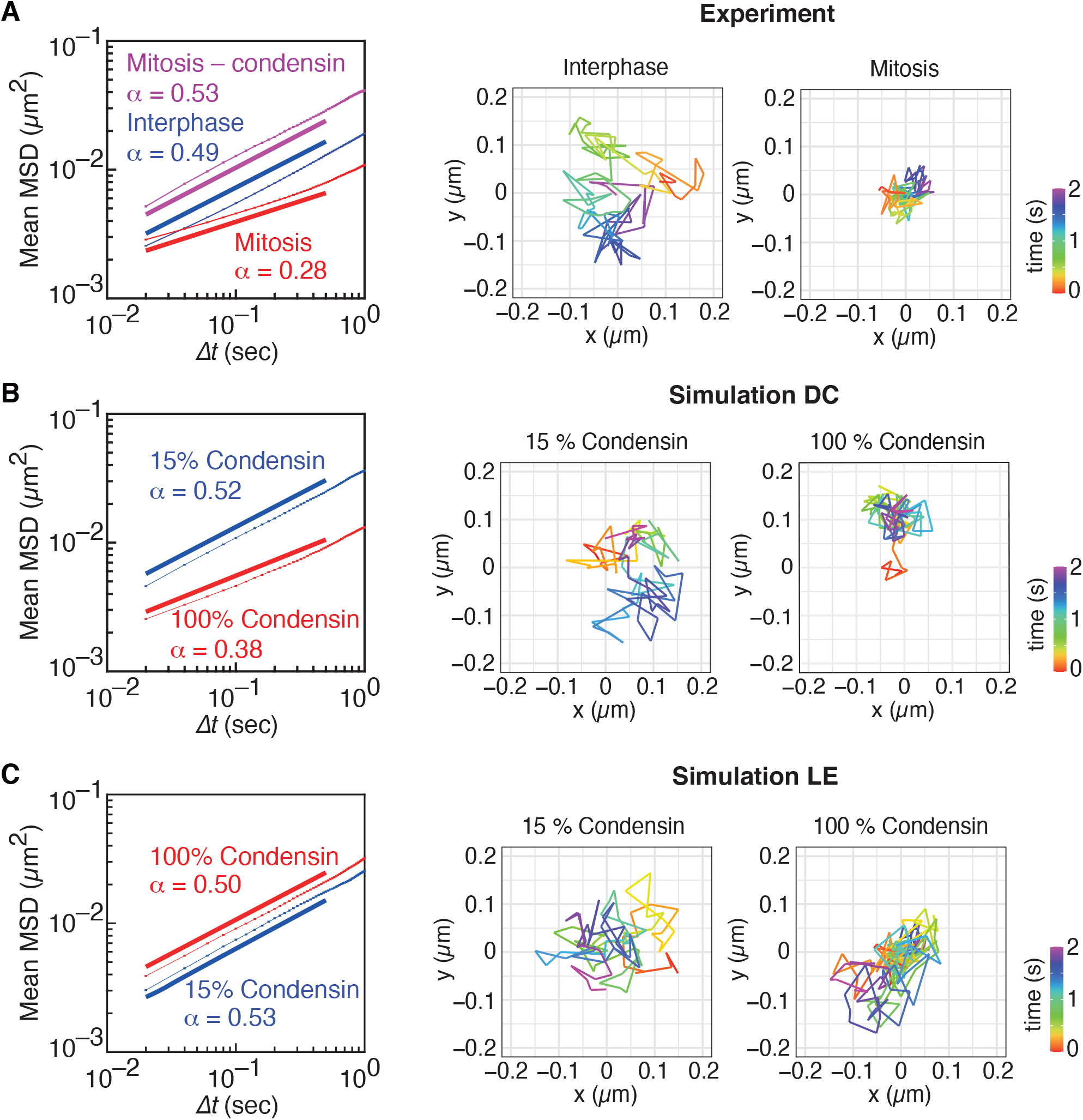
Analysis of mitotic chromatin mobility reduction. **(A)** Experimental MSD analysis of cells in interphase (blue), mitosis (red) and in mitosis following condensin depletion (magenta). 595 interphase trajectories, 271 mitotic control and 149 mitotic without condensin (- condensin) trajectories were analyzed. We calculate an average exponent of power law fits to the MSD of each trajectory up to 0.5 seconds, resulting in the histograms of exponents shown in Supplementary Figure 5A, as well as confidence intervals reported in the text. The solid lines are guide power laws with the respective exponents for comparison. Examples of pseudocolor trajectories in interphase and mitosis during a 2 second window are shown on the right. (**B, C**) *in silico* MSD plots during interphase (blue) or mitotic (red) conditions during simulations of the diffusion capture (**B**) and loop extrusion models (**C**). 2 second traces were analyzed every 60th second during 10 simulation repeats. The mean and confidence intervals are calculated from histograms of exponents to each 2 second trajectory as above, with the mean shown as guide power laws in the plot. Examples of *in silico* bead trajectories are shown on the right.

We next explored the consequences of condensin-dependent *in silico* diffusion capture or loop extrusion on chromatin mobility. Similar to experimental observations, we track chromatin beads in simulation replicates and plot their MSD over time. In the diffusion capture model, the interphase MSD exponent was 0.52 ± 0.03 (mean ± 95% confidence interval, n = 660), close to the experimentally observed value. The exponent was reduced to 0.38 ± 0.02 during *in silico* mitosis. The mitotic mobility reduction is reminiscent of our *in vivo* observations, although the extent of the MSD exponent reduction did not fully reach the experimental observation. An example trajectory of a chromatin bead exemplifies the constrained mitotic mobility due to diffusion capture (Figure 5B). In the loop extrusion model, the interphase MSD exponent was 0.53 ± 0.03. The MSD exponent remained almost unchanged under mitotic conditions when it persisted at 0.50 ± 0.03. A representative bead trajectory further illustrates the largely unchanged mobility (Figure 5C). This suggests that the structural flexibility and dynamics of the chromatin chain is constrained by diffusion capture but remains largely unaltered during loop extrusion.

To study the effects of diffusion capture and loop extrusion on chromatin mobility further, we again turned to condensin titration in our simulations. The mean MSD exponent of the free chromatin chain was 0.53 ± 0.03 consistent with that of an unconstrained Rouse polymer chain with excluded volume (21). Condensin titration in the diffusion capture model sequentially led from an interphase MSD exponent to more and more constrained mobility at full condensin binding site occupancy (Supplementary Figure S5B). The effect of loop extrusion was also condensin concentration-dependent, however did not result in a mean MSD exponent reduction below 0.50 at any of the investigated concentrations. These observations uncover diffusion capture as a powerful mechanism that confines chromatin mobility and that could contribute to the striking mobility reduction observed during mitotic chromosome condensation *in vivo.*

### Mitotic chromatin movements gain anisotropy

In addition to overall constrained mitotic chromatin mobility, expressed in a reduced MSD exponent, we investigated whether mitotic chromosome condensation impacts on the freedom of the directionality of movement, i.e. its anisotropy. We employed an anisotropy metric *η(Δt)* that evaluates whether diffusive movement is equal in x and y directions of the microscope plane, or is constrained in one of the directions more than the other. In effect, *η(Δt)* corresponds to a difference between the diffusion constants in both directions. This metric is most meaningful over short times to probe local directionality constraints. Over longer times, the system locally tumbles resulting in apparent isotropic behavior. For this reason we focus on the average anisotropy *η* over delays of up to 0.1 s.

As a benchmark of our expectations for an isotropic polymer, we first analyzed the anisotropy of our simulated free chromatin chain. We expect *η(Δt) →* 0 as *Δt →* 0, though the finite time resolution of our experiment gives us a finite value for 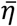. Under our sampling conditions, we find 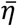 = 0.019 ± 0.005 (Supplementary Figure S6), which means that by random chance we find diffusion in one direction being roughly 75% of that in the orthogonal direction. Applied to our experimental chromatin trajectories, this analysis revealed that chromatin movements in interphase showed greater anisotropy (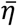 = 0.048 ± 0.005) compared to the isotropic simulated polymer (diffusion in one axis being *64%* of that in the orthogonal direction). The anisotropy became more pronounced in mitosis (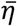 = 0.08 ± 0.01), i.e. movement in one direction was now only 56% of that in the other. This increase in mitotic anisotropy depended on condensin (Figure 6A). It should be noted that the actual experimental anisotropy could be greater, since our microscopy recordings project 3D movements to a random 2D plane, effectively removing any possible difference in diffusivity along the z-axis. We interpret these observations to mean that condensin adds local directional constraint to the diffusive behavior of the chromatin chain in mitosis.

**Figure 6.**
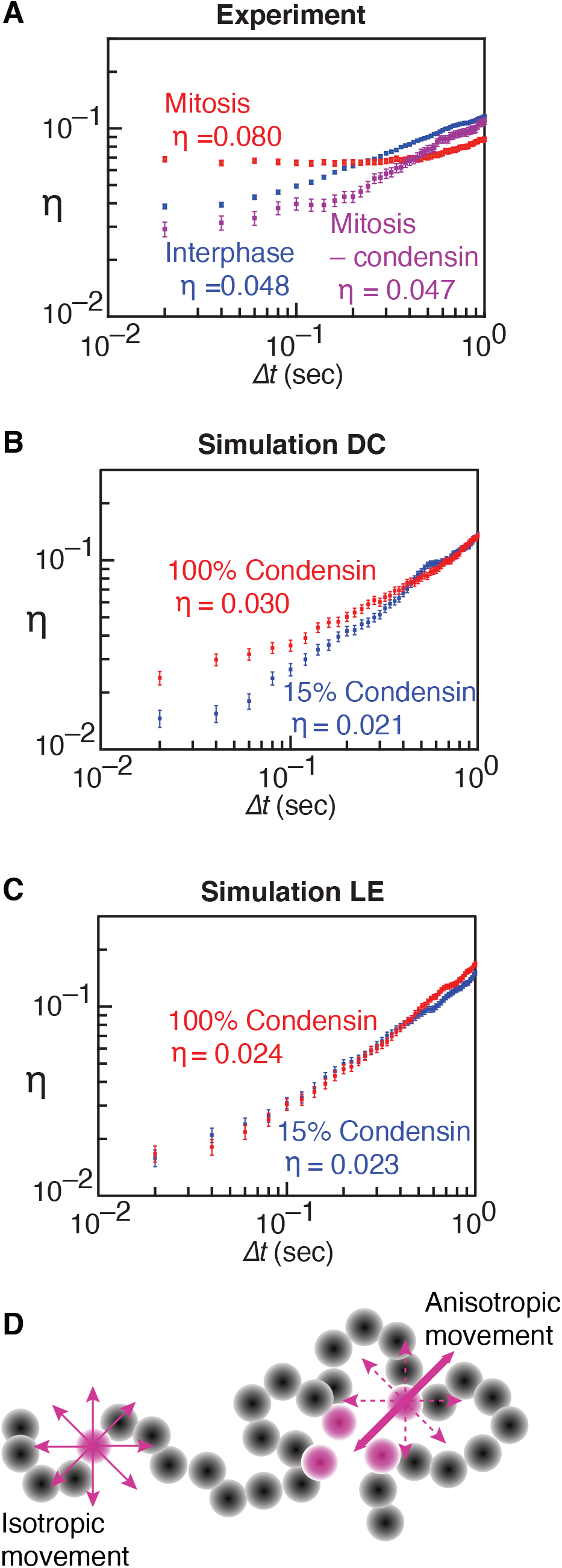
Anisotropy gain of mitotic chromatin movements. **(A-C**) Anisotropy in the chromatin movements used for the MSD measurements in Figure 5 was determined using the *η(Δt)* metric and is shown for the indicated conditions. The means and confidence intervals are shown and were calculated as in Figure 5 over each individual observed or simulated trajectory. (**D**) Schematic for how condensin binding site clustering might introduce directional constraints (anisotropic movement) to the chromatin chain.

We next applied the anisotropy metric to our simulated chromatin movements. Compared to the free chromatin chain, interphase concentrations of condensin slightly increased 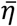 in both the diffusion capture and loop extrusion models (Figures 6B, C). Increasing condensin towards mitotic concentrations barely affected the anisotropy of movements in the loop extrusion model. In contrast, it resulted in a dose-dependent 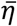 increase in case of the diffusion capture model (Supplementary Figure S6). Qualitatively, therefore, diffusion capture led to increased anisotropy of chromatin movement.

Quantitatively the resultant anisotropy remained below that experimentally observed. We imagine that condensin binding site clustering in the diffusion capture model results in a chromatin topology that constrains chromatin chain movement in certain directions, thus creating local anisotropy that we can experimentally and computationally detect (Figure 6D). The effect might be more pronounced *in vivo* where additional chromosome-bound proteins might augment any constraints. The overall more dynamic nature of the chromatin chain in the loop extrusion model did not create a similar phenomenon.

### Condensin cluster formation within mitotic chromosomes

Given the above contrasting observations of how diffusion capture and loop extrusion impact on mitotic chromosome behavior, we wanted to directly visualize the emergent 3D organization of condensin inside mitotic chromosomes. To this end, we performed stochastic optical reconstruction microscopy (STORM) to visualize condensin within fission yeast mitotic chromosomes at high spatial resolution. We arrested fission yeast cells in mitosis by transcriptional repression of the Slp1 activator of the anaphase promoting complex (26). Condensin’s Cut14 subunit was fused to a SNAP tag, which we labeled with an Alexa Fluor 647 dye following cell fixation and permeabilization. STORM imaging now allowed us to determine the location of condensin molecules within the fission yeast nucleus. The particle count per nucleus was 1114 ± 110 (median ± S.E.M, n = 19), in line with the expected number of condensin molecules (41,42). Qualitatively, condensin molecules appear to cluster in small groups that are widely scattered throughout the nucleus (Figure 7A; a partial volume corresponding to the chromosome I left arm is depicted in Figure 7B). To quantitatively describe condensin clustering, we performed DBSCAN cluster analysis of the condensin distribution (see Materials and methods). This revealed a predominance of small clusters with 2 to 4 condensin molecules while larger clusters with 10 or more condensins were also detected, but less frequently (Figure 7C).

**Figure 7.**
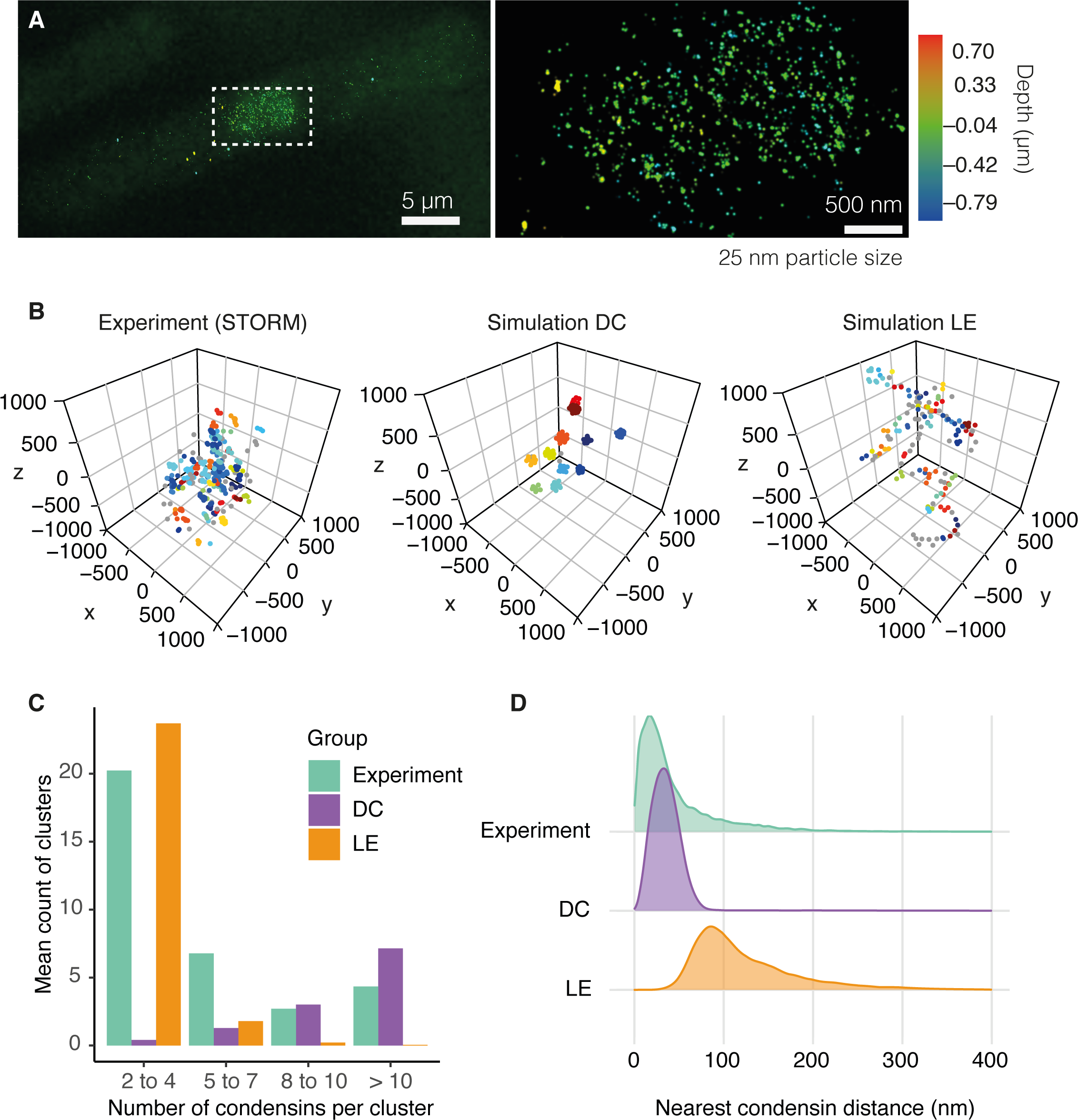
Patterns of condensin distribution in 3D space. (**A**) STORM image of condensin in a mitotic fission yeast cell, a magnified view of the nuclear area is shown on the right. Detected particles are shown as dots with 25 nm diameter, their pseudocolor represents the depth within the nucleus. (**B**) Spatial distribution of condensin from a region of similar size as the chromosome I left arm seen in the experiment (left), as well as condensin distributions in representative conformations from mitotic diffusion capture (middle) and loop extrusion (right) simulations. Particle clustering used a 100 nm threshold and a minimum particle number per cluster of 2. Distinct colors reflect distinct clusters. Condensins not contained in a cluster are shown in grey. (**C**) Cluster size distribution in the experimental and simulated condensin localizations. Mean cluster counts from 19 images, and from 1,200 snapshots at 10 second intervals throughout 10 simulation replicates are shown. (**D**) Nearest condensin distance for each condensin as a quantitative descriptor of condensin clustering in the STORM experiment (green), diffusion capture (purple) and loop extrusion (orange) simulations.

We next performed a similar analysis of the condensin distribution in our simulated mitotic chromosomes formed by diffusion capture or loop extrusion. Qualitatively, diffusion capture led to the formation of condensin clusters of various sizes, spread throughout the chromosome volume. Loop extrusion, in contrast, resulted in the formation of an apparent condensin backbone, consisting of approximately evenly spaced condensin molecules (Figure 7B). When we subject these condensin distributions to the same quantitative cluster analysis, we find that diffusion capture results in a broad distribution of cluster sizes, skewed towards large clusters. In contrast, rarely more than two condensin molecules were found to cluster during loop extrusion (Figure 7C). Both simulated distributions differ from the experimental result. While the experimental result and the diffusion capture model contain a range of cluster sizes, the median cluster size in the diffusion capture simulations was 12, which is distinctly larger than the experimentally observed median cluster size of 4. If condensin clusters form by diffusion capture *in vivo,* a mechanism must exist that limits their size.

As another quantitative metric to compare the condensin distributions within native and simulated chromosomes, we plotted the distances of each condensin molecule from its nearest neighbor. The condensin clustering observed in the experimental data, as well as in the diffusion capture simulations, mean that the majority of condensins possess a close neighbor. The median nearest distance was 29 nm in our STORM data and 34 nm in the diffusion capture simulations (Figure 7D). Condensins in the loop extrusion model were spread out along the chromosome backbone with a greater median distance from their nearest neighbors of 107 nm. This analysis confirms a clustering pattern that is generated by the diffusion capture mechanism that is lacking from the loop extrusion simulations.

## DISCUSSION

We computationally examined the consequences on chromosome formation of two prevalent models of condensin function, loop extrusion and diffusion capture. At their essence, both models result in the establishment of loops between distant sites along a chromatin chain. Only the mechanisms by which these loops form differ. In the case of loop extrusion, active movement of the chromatin chain results in loop growth. Diffusion capture, in contrast, takes advantage of stochastic loop formation by Brownian motion. Condensin in the latter case merely acts to stabilize such loops for a period of time. These parallels and distinctions result in similarities between chromosomes that form by both mechanisms, but also in a number of differences.

### Implications for chromosome dimensions and chromatin density

As a consequence of loop formation, both loop extrusion and diffusion capture can recapitulate experimentally observed chromatin contact distributions that develop during mitotic chromosome formation. Chromatin loops, created by either loop extrusion or diffusion capture, furthermore can result in chromosome axis shortening. While diffusion capture results in robust and dose-dependent chromosome compaction, loop extrusion displays a more complex relationship between the number of loop extruding condensins and the resultant chromosome dimensions.

In our simulations, we assume that one condensin is active per every approximately 20 kb of chromatin. This estimate stems from experimentally observed condensin ChIP distributions (22,27,34) as well as quantitative estimates of roughly 1000 condensin complexes per fission yeast cell nucleus (41,42). During our mitotic loop extrusion simulations, all these complexes are equally active in extruding loops and in initiating nested loops, based on simple probability. This results in a chromosome backbone that is longer than experimentally observed and, notably, is longer in mitosis than in interphase. To achieve native-like chromosome axis shortening by loop extrusion, it is possible that additional levels of condensin regulation tune loop intervals and loop nesting. While such mechanisms remain to be explored in organisms that rely on a single source of condensin, like fission yeast, the existence of two distinct condensins in other organisms could facilitate such regulation.

The condensin density on human chromosomes is similar to that in fission yeast (approximately 1 condensin per 20 kb (43)). If human condensin shapes chromosomes by loop extrusion, we should expect loop sizes and chromosome dimensions to be sensitive to changes in condensin concentration. Against this expectation, chromosome shape is remarkably insensitive to substantial reductions in condensin levels (44,45). It will be interesting to study the consequences of altered condensin concentrations on simulated human chromosome formation (10), as well as on chromosome condensation in a defined experimental system (46).

### Implications for chromatin mobility

Mitotic chromosomes are not a static end-product of chromosome condensation, they are dynamic entities whose integrity is maintained through continued condensin ATP hydrolysis cycles (47). In the loop extrusion and diffusion capture models, continued ATP hydrolysis maintains chromosome architecture in different ways that make distinct testable predictions about chromosome properties. In the diffusion capture model, condensin dissociation and reassociation gives condensin clusters the plasticity to evolve by merging or splitting. The net consequence of condensin clustering, however, is to limit chromatin movements and to impose anisotropy. This is borne out in our experimental observations that revealed both a reduced MSD exponent as well as increased anisotropy. In contrast, the loop extrusion model envisions that dissociating condensins initiate new loops that grow again by directional enlargement. In our simulations, loop extruding condensins turn over on average every 2 minutes. While condensin turnover on mitotic fission yeast chromosomes remains to be measured, 2 minutes corresponds to relatively stable association, when compared to budding yeast condensin or even human condensin II (48–50). Despite the therefore relatively slow turnover of condensin in our loop extrusion model, chromatin remains mobile and unconstrained in the directionality of movements, contradicting our experimental observations.

While chromatin movements in interphase are close to what is expected from an unconstrained Rouse polymer with excluded volume (21), our experiments point to a markedly smaller MSD exponent in mitosis, potentially as small as 0.23. This reduction is partly reproduced by the diffusion capture model. An exponent of ¼ has been described to arise from the behavior of long ring polymers in a melt or from ring polymers in a set of fixed obstacles (51,52), suggestive of a potential role of chromatin loops in the sub-diffusive behavior observed in mitosis. Understanding the quantitative nature of the observed diffusive behavior, in light of biophysical models of chromosome formation, remains an open challenge.

### Condensin cluster formation within mitotic chromosomes

A predicted feature from the diffusion capture model is the formation of condensin clusters of variable sizes, spread throughout chromosomes. The loop extrusion model, in contrast, predicts that condensins are spaced out along a chromosome backbone. We could not discern such a chromosome backbone in our STORM images of mitotic fission yeast cells. Rather, condensin was found in dispersed small foci. While these foci are reminiscent of those predicted by diffusion capture, their median cluster size was smaller than observed in our simulations. We note that cluster size in our simulations is principally restricted by steric constraints created by the chromatin chain. These steric constraints can be expected to be greater *in vivo*, where numerous proteins in addition to histones decorate the chromatin chain. Such additional constraints offer one possible explanation for why cluster sizes might be smaller *in vivo.* Alternatively, other properties of condensin or of its chromosomal binding sites might limit the size of the clusters that can form.

High resolution imaging of condensin in human chromosomes, using stimulated emission depletion (STED) microscopy, has also revealed condensin clusters instead of a continuous condensin backbone (43). While appearing overall scattered, these clusters were enriched towards axial positions inside human chromosomes. Loop extrusion is a powerful mechanism to explain axial enrichment. Expanding loops move outwards while pushing loop anchors towards the center. Could it therefore be that condensin shapes human chromosomes by a combination of diffusion capture and loop extrusion? To achieve loop extrusion, condensin has been proposed to employ an intrinsic motor, as observed *in vitro*(4,7). However, it remains uncertain whether condensin can extrude densely packed chromatin loops *in vivo.* We therefore suggested that loops that are established by diffusion capture could expand by means of an extrinsic motor, e.g. RNA polymerases that are known to reposition condensin along transcription units (34,53,54). Such an extrinsic ‘loop expansion’ mechanism (2) could similarly result in axial condensin cluster accumulation. To further explore this question, it will be interesting to analyze the condensin distribution in chromosomes that form in the absence of transcription (55).

### Outlook

A perceived benefit of loop extrusion is that it provides a fool-proof mechanism to ensure that condensin-dependent chromatin interactions happen within the same chromatin chain, rather than between neighboring chromosomes. However, experimental observations suggest that condensin promotes interactions both within as well as between chromosomes (16–18). If diffusion capture is blind as to whether interactions are established within or between chromosomes, how can we explain condensin’s ability to individualize chromosomes? Even diffusive interactions are always more likely to occur within a continuous chromatin chain, as compared to an interaction with an independently moving chromatid or chromosome. This provides an inherent mechanism that sufficiently explains chain compaction and individualization of small budding yeast chromosomes (11). We suggest that, in larger chromosomes, loop expansion following diffusion capture aids chromosome individualization as outwards moving loops repel each other.

Lastly, our diffusion capture model is a specific case of a string-and-binder polymer model (56,57). A feature of such models is that they can lead to a collapse of the polymer chain into a dense ball. We found this to be the case only when condensin binding sites were much more closely spaced than experimentally observed. Using actual condensin spacing, local clusters form that are isolated from neighboring hubs by steric constraints. While clusters evolve over time by dynamic exchange of condensin binding sites, the overall chromosome remains in a stable steady state. We have provided arguments to suggest that diffusion capture can make an important contribution to mitotic chromosome formation. In how far this mechanism cooperates with intrinsic loop extrusion, or with extrinsic loop expansion, to shape chromosomes remains a fascinating question to address by further integrative computational and experimental studies.

## AUTHOR CONTRIBUTIONS

T.G., X.F., Y.K., P.A.B. and F.U. together conceived and developed the study. T.G. and X.F. wrote the computer code, performed simulations and formatted data to compare with the experiments. Y.K. performed all the biological experiments, B.K. developed the MSD and anisotropy analyses, C.B. analyzed Hi-C data, T.G., P.A.B. and F.U. wrote the manuscript with the input of all co-authors.

## ACKNOWLEDGEMENT

We would like to thank D. Aubyn in the Crick Advanced Light Microscopy Science Technology Platform for her help with STORM, D.J. Barry and M. Way for help with particle tracking, A. Rabinowitz for Hi-C data analyses, the Crick Advanced Sequencing Science Technology Platform, M. Molodtsov and all our laboratory members for valuable discussions and comments on the manuscript.

## FUNDING

This work was supported by the European Research Council under the European Union’s Horizon 2020 research and innovation program (grant agreement No 670412), The Francis Crick Institute, which receives its core funding from Cancer Research UK (FC001003, FC001198), the UK Medical Research Council (FC001003, FC001198), and the Wellcome Trust (FC001003, FC001198), as well as the Japanese Society for the Promotion of Science (JSPS) and Waseda University (grant for special research projects 2020C-738) to Y.K. Funding for open access charge: The Francis Crick Institute.

## CONFLICT OF INTEREST

The authors declare no conflict of interest.

## Supplementary Data

**Supplementary Figure 1.**
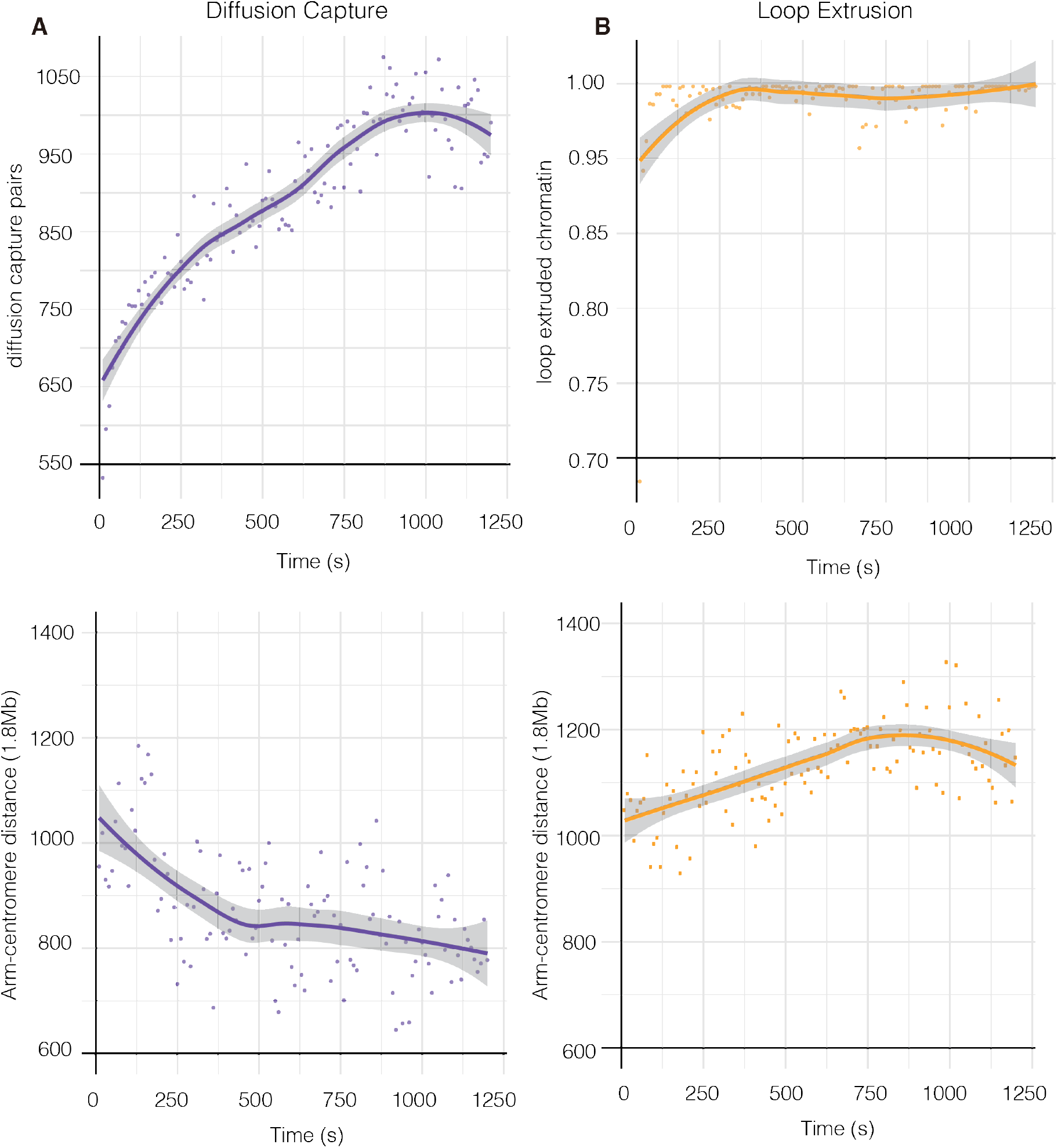
Time development of mitotic diffusion capture and loop extrusion simulations. (**A**) Diffusion capture simulations. Starting from a relaxed chromatin chain, the time development of diffusion capture pair formation, as well as the axial chromosome distance of the *in silico* fluorophore pair analyzed in Figure 2, are plotted over time. The medians from the 10 simulation repeats are shown at 10 second intervals. A smoothened fit and its 95% confidence interval are also shown. (**B**) Loop extrusion simulations. As in (**A**), but the percentage of the chromatin chain that is contained in loops, as well as the *in silico* fluorophore pair distance, are shown.

**Supplementary Figure 2.**
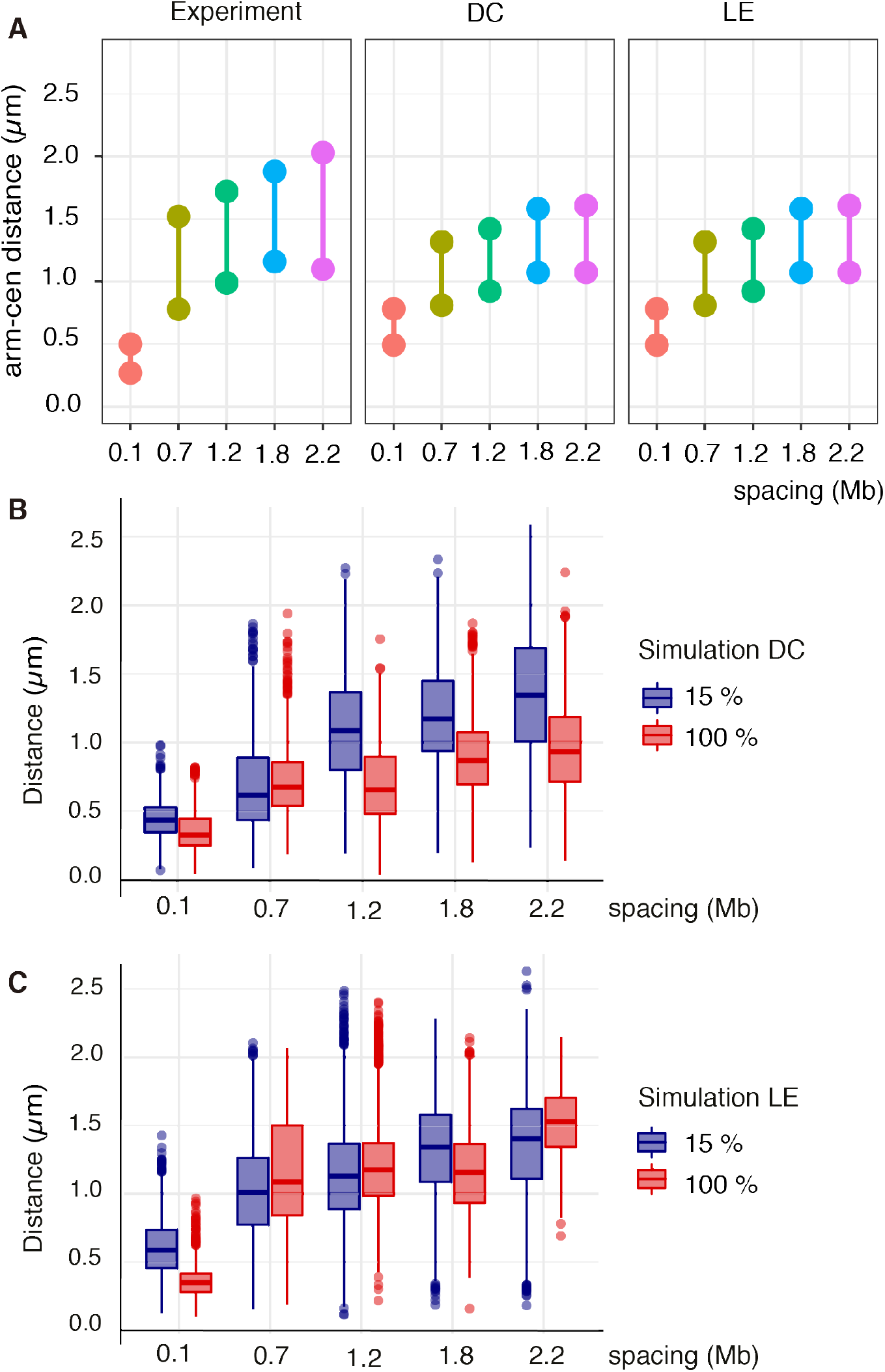
Axial chromosome lengths in experimental data and diffusion capture and loop extrusion simulations, measured over different genomic distances. (**A**) Line plots depict minimal and maximal values of distances recorded of fluorophore pairs at the indicated genomic spacing during experimental interphase (left panel, data from reference (26)), as well as the interquartile ranges from 1,200 measurements at 10 second intervals during 10 simulation repeats during diffusion capture (middle panel) and loop extrusion (right panel) simulations using interphase condensin levels. (**B, C**) Euclidean distance distributions of *in silico* fluorophores at the indicated genomic spacing for diffusion capture (**B**) or loop extrusion simulations (**C**), using interphase (blue, 15% condensin) and mitotic conditions (red, 100% condensin). Boxes depict medians and interquartile ranges.

**Supplementary Figure 3.**
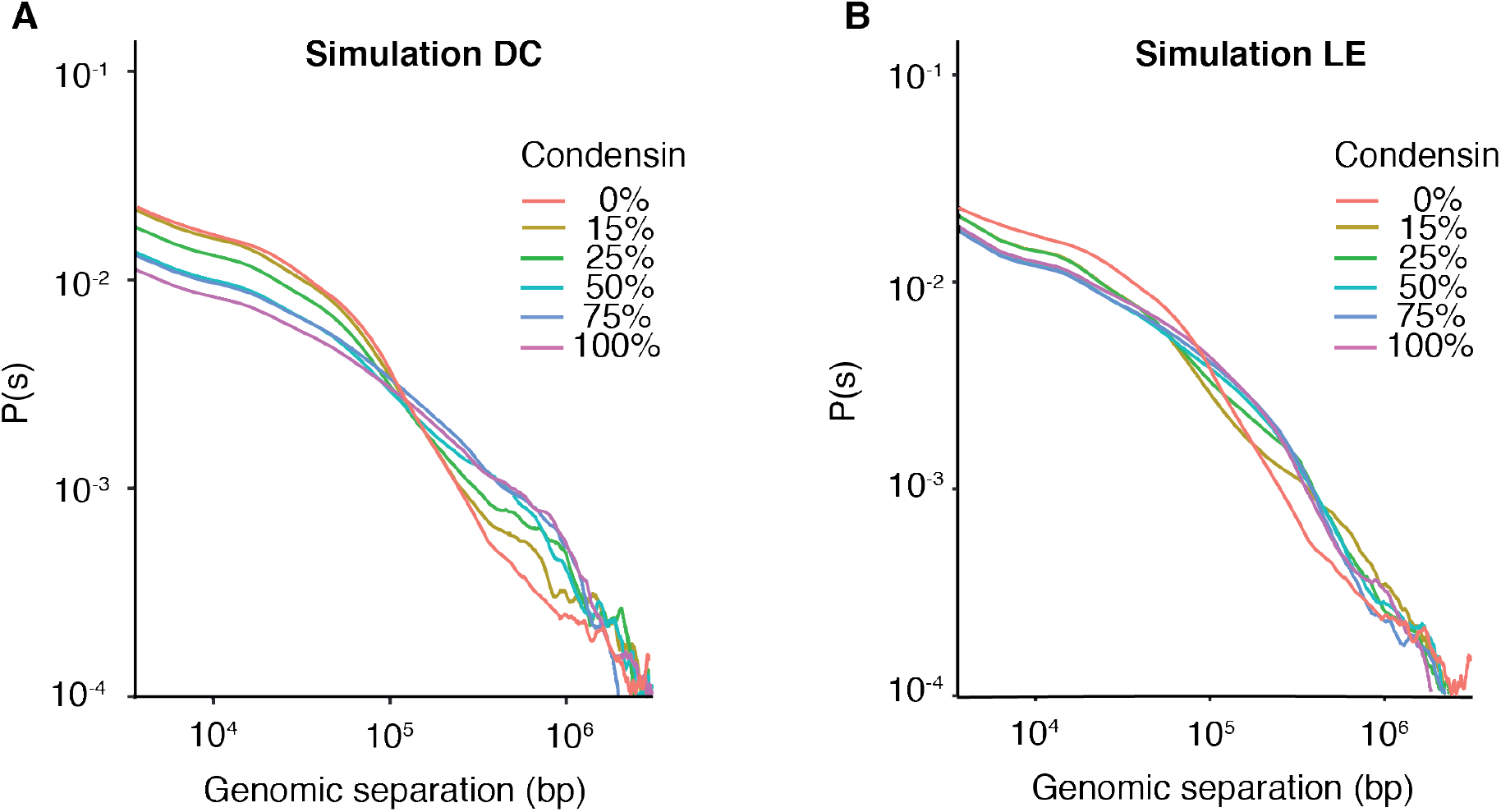
Condensin concentration-dependent changes to the contact probability distribution in the diffusion capture and loop extrusion models. (**A, B**) Contact probability as a function of genomic separation, as in Figure 3, is plotted for different condensin concentrations during diffusion capture (**A**) and loop extrusion simulations (**B**). 12,000 conformations, recorded at 1 second intervals during 10 simulation replicates, were analyzed in each case.

**Supplementary Figure 4.**
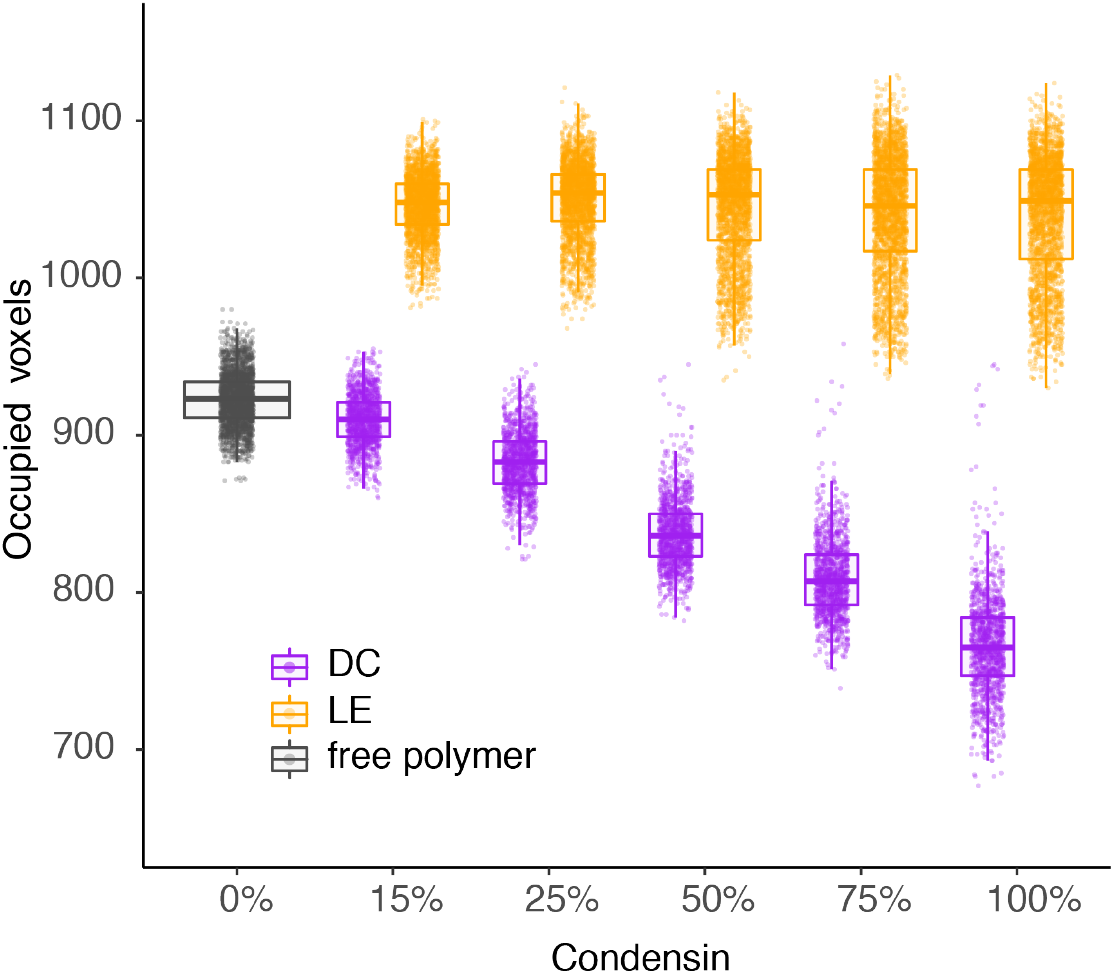
Condensin concentration-dependent chromosome volume compaction during simulated diffusion capture and loop extrusion. Occupied voxel distributions during diffusion capture (purple) and loop extrusion (orange) at different condensin concentrations is shown. 1,200 snapshots, taken at 10 second intervals from 10 simulation replicates were analyzed. Boxes indicate the medians and interquartile ranges.

**Supplementary Figure 5.**
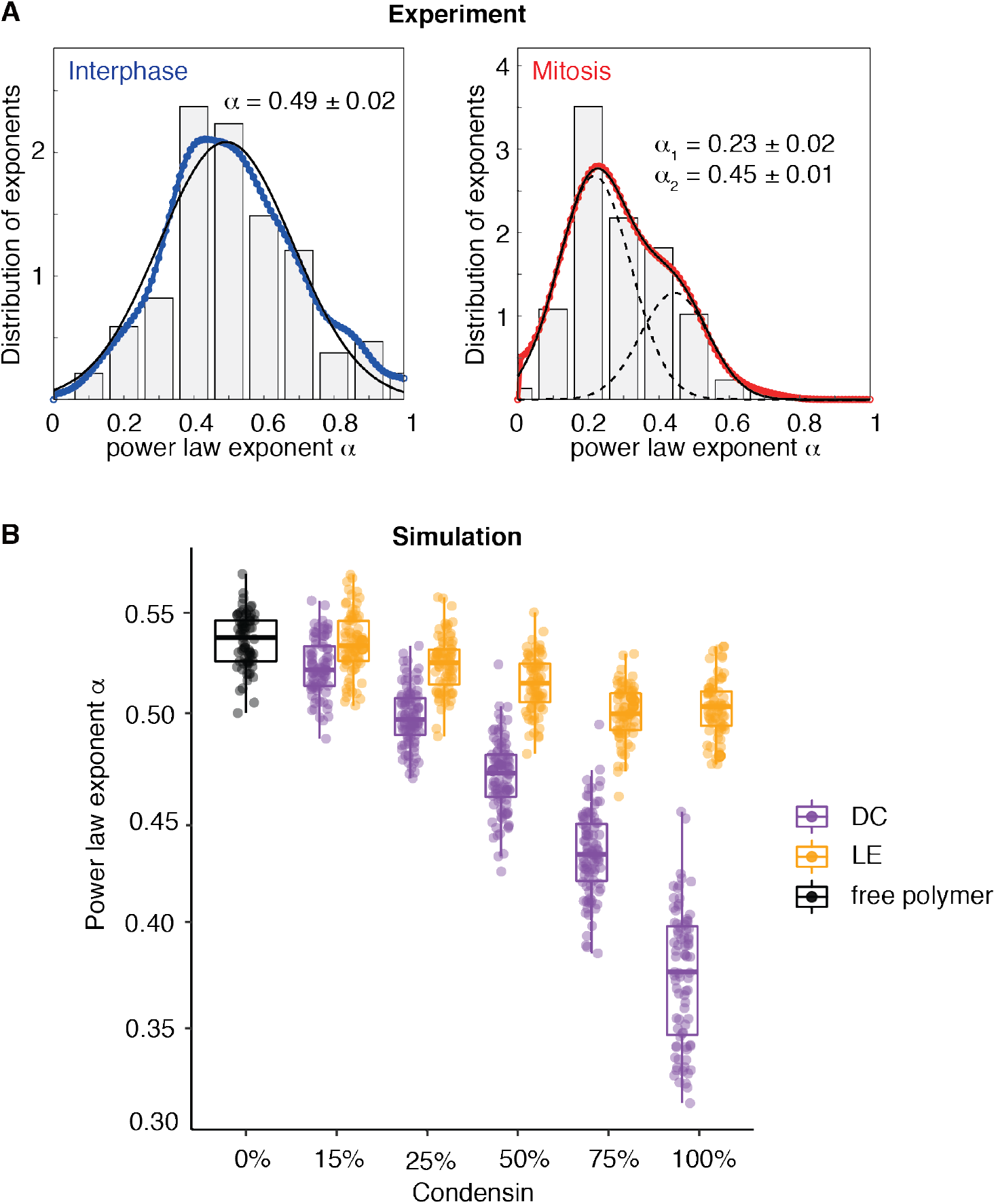
Additional analyses of *in vivo* and *in silico* chromatin mobility. (**A**) Histogram (grey bars) and kernel density estimate (colored circles) of probability density of individual MSD exponents in interphase and mitosis. The solid black line represents a single Gaussian fit of the interphase distributions (n = 595), while the dashed lines show a double Gaussian fit for the exponent distribution in mitosis (n = 271). (**B**) Condensin concentration-dependence of chromatin chain mobility in the diffusion capture and loop extrusion models. The distribution of MSD exponents of a free chromatin chain (black) is compared to the indicated condensin concentrations during diffusion capture (purple) and loop extrusion simulations (orange). 2 second traces were analyzed every 60th second during 10 simulation repeats. Boxes indicate the medians and interquartile ranges. We previously used similar simulations to arrive at an MSD exponent of a free polymer chain, α = 0.57 ± 0.11 (reference 22; here 0.53 ± 0.03) compatible within the confidence intervals of both studies. A difference in approach was the use of a nuclear constraint in our current study, which was not included in our previous simulations and which can be expected to impose a limit on mobility.

**Supplementary Figure 6.**
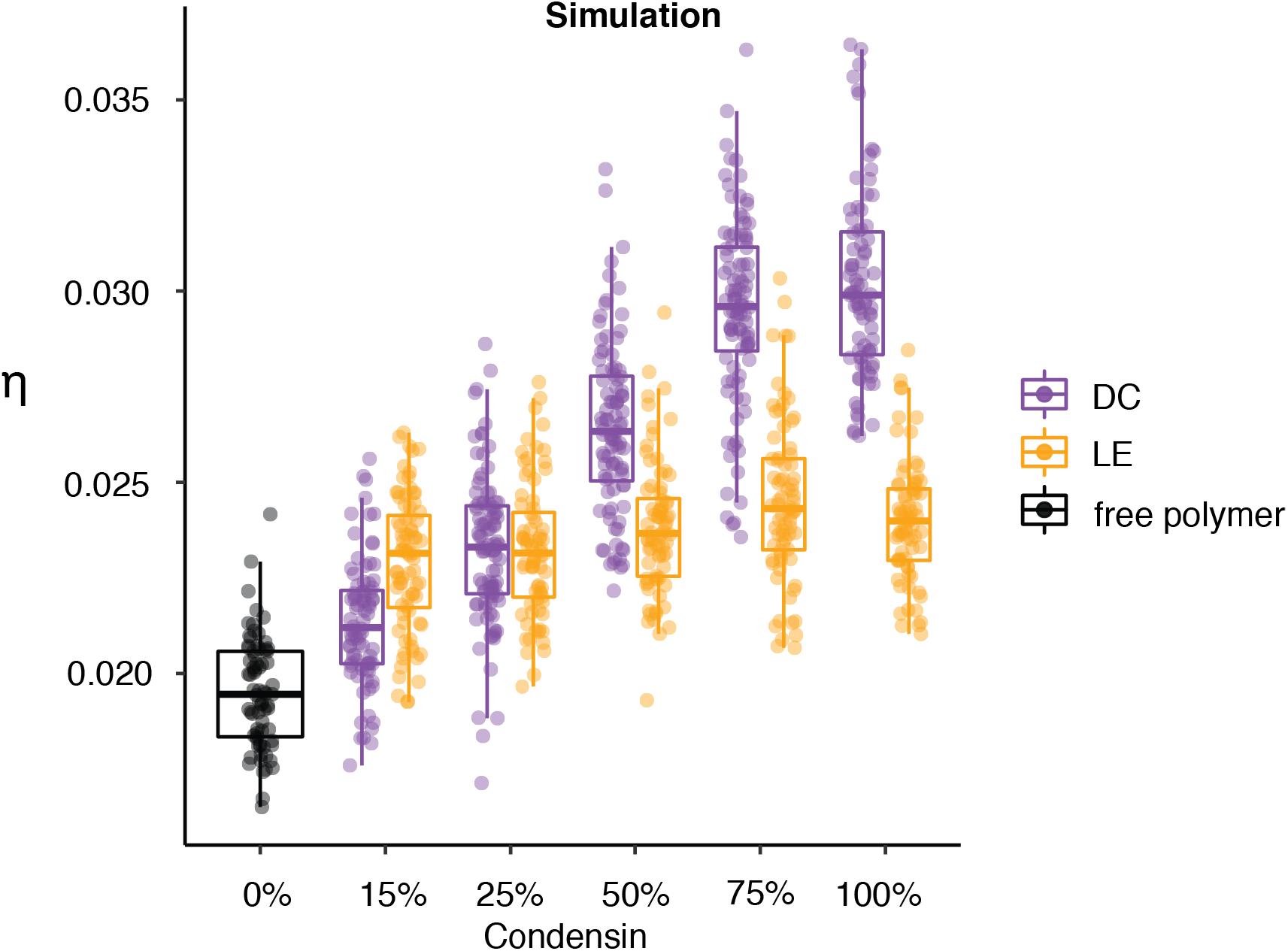
Condensin concentration-dependence of anisotropic motion during diffusion capture and loop extrusion. The distribution of anisotropy exponents of a free chromatin chain (black) is compared to the effect of the indicated condensin concentrations during diffusion capture (purple) and loop extrusion simulations (orange) shown in Supplementary Figure S5. Boxes indicate the medians and interquartile ranges.

**Supplementary Table S1.**
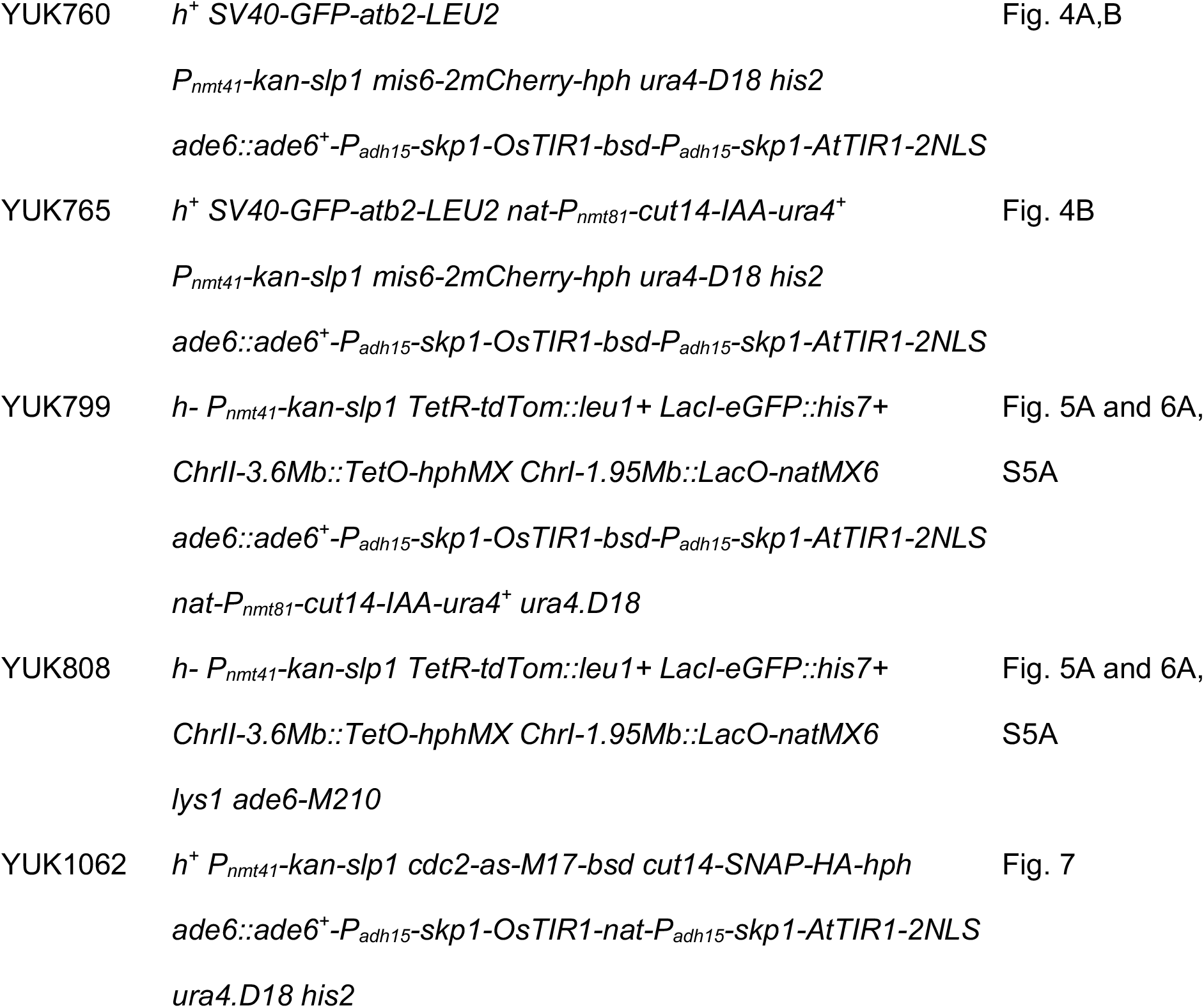
Yeast strains used in this study.

## Notes

### Competing Interest Statement

The authors have declared no competing interest.

